# Iterative Extracellular Vesicle Protein Co-Expression Biomarker Refinement for Preoperative Classification of Histopathological Growth Patterns in Colorectal Liver Metastasis Patients

**DOI:** 10.64898/2026.02.02.702621

**Authors:** Rosalie Martel, Molly L. Shen, Migmar Tsamchoe, Stephanie K. Petrillo, Anthoula Lazaris, Peter Metrakos, David Juncker

## Abstract

Preoperative triage of colorectal liver metastases (CRCLM) by histopathological growth pattern (HGP)—angiogenesis-dependent desmoplastic (dHGP) and vessel co-opting replacement (rHGP)—could guide anti-angiogenic therapy, yet HGP scoring requires resected tissue. We present an extracellular vesicle (EV) inner and outer protein (EVPio) co-expression assay and iterative biomarker refinement for plasma-based HGP classification. We established a minimal, high-throughput plasma pre-processing workflow (low-speed centrifugation and 0.45 μm filtration) with comparable EVPio assay performance to size-exclusion chromatography. We established an EV biomarker selection template with growing cohorts—feasibility (*n* = 3), pilot (*n* = 9), discovery (*n* = 67)—ranking candidate protein pairs by signal quality (SNR, CV), redundancy (inter-correlation/orthogonality), and HGP separation (effect size, significance, ROC). This process reduced an initial 19×18 capture/detection set to a focused 9×9 panel (81 co-expression pairs). In a 58-patient CRCLM subset (22 dHGP, 14 rHGP, 22 mixed), three pairs achieved high signal quality with significant differential expression across HGPs. A three-feature linear discriminant model yielded 75.9% cross-validated accuracy (AUC 0.77) for classifying pure dHGP vs. non-dHGP. The results show that co-expression signatures capture defining features of HGP biology while revealing heterogeneity. The proposed EV biomarker refinement template is generalizable and our results show that co-expression signatures capture defining features of HGP biology supporting efforts towards clinically actionable, HGP-driven therapeutic guidance.

---

Colorectal cancer (CRC) is the third most diagnosed cancer and second leading cause of cancer death worldwide.^1^ While its incidence in older adults has steadily decreased in most high-income countries, the last few decades have seen an uptake of cases in younger adults, with ties to widespread changes in diet and lifestyle factors.^2,3^ The proportion of advance-stage disease (with regional or distant spread) at diagnosis has also been increasing in the past 10-15 years, reaching 60% in 2019 in the United States.^4^ Overall, it is estimated that 25%-30% of all CRC patients will develop metastases, primarily to the liver.^3,5^

Colorectal cancer liver metastases (CRCLM) can present with distinct histopathological growth patterns (HGPs) that are defined by their interaction with the surrounding liver tissue and how they access blood supply. Two of the most commons HGPs are the desmoplastic (dHGP) and replacement (rHGP) presentations. As their names suggest, dHGP tumors isolate themselves from the local liver parenchyma with a desmoplastic rim, while the rHGP tumors effectively “replace” the existing tissue by infiltrating into the adjacent liver parenchyma.^6,7^

Underlying these structural differences are two separate strategies to secure tumoral blood supply: dHGP metastases rely on angiogenesis, while rHGP metastases forego de novo vessel production in favor of co-opting existing liver blood vessels.^6,7^ These divergent behaviors have important therapeutic implications: anti-angiogenic agents such as the anti-VEGF therapy bevacizumab (Bev) are less likely to be effective in rHGP tumors compared to dHGP tumors. Desmoplastic tumors receiving neoadjuvant Bev and chemotherapy have more than double the 5-year overall survival compared to patients with rHGP lesions who have received the same neoadjuvant regimen.^8–10^ Currently, HGP typing requires histological analysis of resected samples obtained during surgical resection and is thus seldom available to guide systemic therapy selection.^6^

Liquid biopsies have emerged as a less invasive alternative to capture the spatial and temporal heterogeneity of tumors by sampling the biological materials dynamically released by cancer cells in the systemic circulation.^11,12^ Amongst the candidate liquid biopsy analytes, extracellular vesicles (EVs), a collection of small, cell-derived membrane bodies involved in intercellular communication,^13,14^ have several attractive characteristics: a high abundance in biological fluids, compatibility with -80°C storage, and a complex cargo of interrelated biomolecules (proteins, nucleic acids, lipids).^15^ Owing to their dynamic expression and active participation in biological pathways, EV proteins are particularly well-suited analysis targets for clinical applications.^16^ Indeed, an increasing number of EV protein-based platforms have shown promise in detecting, differentiating, staging and monitoring various cancers.^17–22^

We previously developed the Extracellular Vesicle Antibody Microarray for Multiplexed Inner and Outer Protein Analysis (EVPio) to probe pairwise antigen co-expression and perform comparative protein phenotyping in CRC cancer cell line-derived EVs.^23^ EVPio incorporates custom antibody microarrays for high-throughput analysis, an antigen retrieval (AR) treatment to enable intravesicular probing, and barcoded antibodies with complementary fluorescent branched DNA trees for multiplexing and signal amplification.^23^ Here, we introduce a dedicated EVPio-CRCLM assay for HGP-based classification (**Figure 1**). To this end, we (i) optimized a minimal sample processing step upstream of the EVPio assay to enable plasma EV analysis (**Figure 1A**); (ii) developed the EVPio-CRCLM assay iteratively, including its protein biomarker panel and experimental parameters, using clinical insights and pilot phenotyping data; and (iii) combined EVPio-CRCLM phenotyping and LDA-based statistical modelling to perform the final HGP classification in a cohort of 58 CRCLM patients (**Figure 1B**).

**Figure 1.**
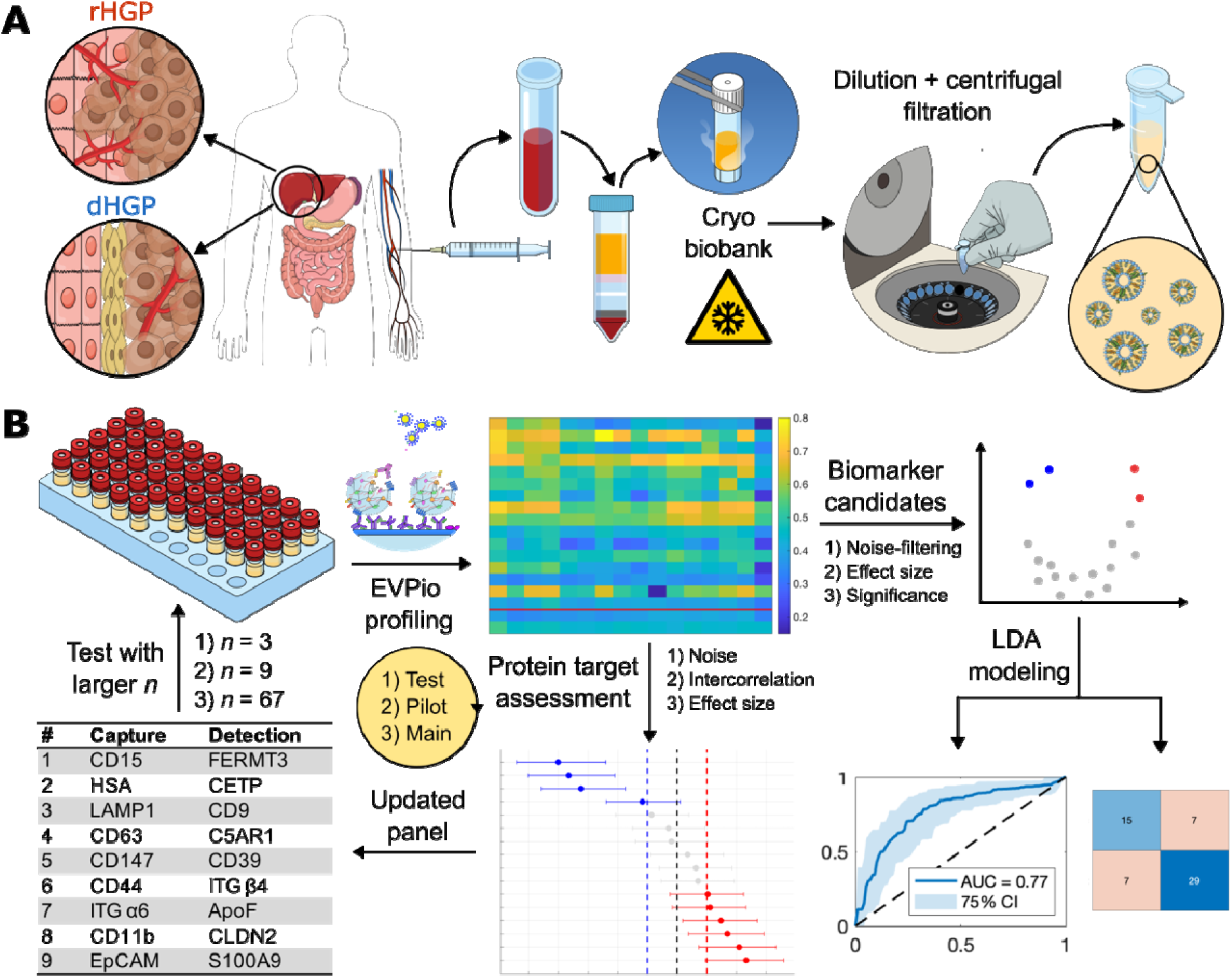
EVPio-CRCLM for HGP-based classification. (**A**) Biobanked plasma samples collected from CRCLM patients with rHGP or dHGP liver metastases underwent minimal processing (dilution in PBS, centrifugal filtration) prior to EVPio phenotyping. (**B**) The EVPio-CRCLM assay was iteratively refined through successive EV protein co-expression profiling of progressively larger CRCLM plasma sample cohorts (number of included patients *n*). Biomarker panel optimization was guide by analytical insights such as biomarker quality, intercorrelation and HGP differentiation effect size. This figure includes elements adapted from ScienceFigures.org, Open Design License 1.1 (see **Table S4**)

## Results

### EVPio supports plasma EV analysis following minimal sample processing

With features like superior vesicle purity and integrity, modest equipment requirements (e.g., no ultracentrifuge required) and a shorter processing time than other EV separation techniques (∼20 min),^24^ size exclusion-chromatography (SEC) was a natural choice to isolate EVs for cell line EVPio.^23^ However, when working with numerous low-volume plasma samples, SEC drawbacks such as limited parallelization potential, dilution effects and lower yield (necessitating higher sample volumes) become more constraining.^24^ Plasma contains a systemic mix of EVs from multiple systems and organs, and biomarker-bearing, disease-relevant EVs can be rare and difficult to detect.^25^ Dilution and losses during isolation further exacerbate these challenges. For example, dyed, purified cancer cell line (HT29) EVs spiked 1/8 in phosphate buffer saline (PBS) and captured on antibody microarrays had a median 4.4-fold reduction in detection signal-to-noise ratio (SNR) if isolated with SEC prior to capture. In the presence of a plasma matrix, which appears to partly protect against SEC-mediated signal loss at higher dilutions, the median SNR is reduced 1.9-fold for 1/8 EV spike-ins (**Figure S1**).

To adapt EVPio to the analysis of plasma EVs, we aimed to develop a high throughput, streamlined sample processing pipeline that leverages EVPio assay specifics to maximize efficiency. Since EVPio relies on two distinct antigen recognition steps—a capture step to pull down specific subpopulations, and a detection step to probe a given target within that subpopulation—free proteins are intrinsically excluded from detection signals, and the main source of extraneous signal are larger impurities like cell membrane fragments. Consequently, we trialed combinations of low-speed centrifugation, concentration and ultrafiltration, varying the filter pore size and concentration factor. Beyond a simplified and easily scalable workflow, a filtration-based approach has the added benefit of retaining small (<50 nm) EVs, which are often lost in many established purification methods.^26^

Conditions were evaluated based on the SNRs obtained in a model EVPio assay of nine capture and three detection targets carried out on serially diluted processed samples (**Figure S2**). For the ultrafiltration step, we compared the subpopulation SNR distributions obtained with pore sizes of 0.22 μm and 0.45 μm due to their widespread use in EV sample preparation^27^ (**Figure 2A**). SNRs from the 0.45 μm filtration conditions were 2.3× higher than with 0.22 μm filtration, likely due to the signal contributions of larger EVs excluded from 0.22 μm-filtered samples. Interestingly, the improvement in SNR varies by capture target, with EGFR^+^, ITG β1^+^ and ITG α2^+^ EVs showing the sharpest increases. Such subpopulations may contain a higher proportion of larger EVs and suffer lower detection sensitivity with decreasing filtration cutoffs. Plasma pre-concentration was also tested to see if it could boost the signal of low-abundance antigen pairs but was found to be of limited utility owing to modest SNR improvements and experimental complications due to increased viscosity (**Figure S2**). Benchmarked against SEC isolation (**Figure S3**), 0.45 μm filtration-based plasma sample processing yielded overall comparable SNRs (median ratio of 0.8 compared to SEC), with some subpopulation- and dilution factor-specific variation (**Figure 2B**). The higher EVPio SNRs of some phenotypes with SEC isolation, e.g., EGFR^+^ and EpCAM^+^ EVs, may be explained by a high protein target abundance in the tighter size range enriched within retained fractions.

**Figure 2.**
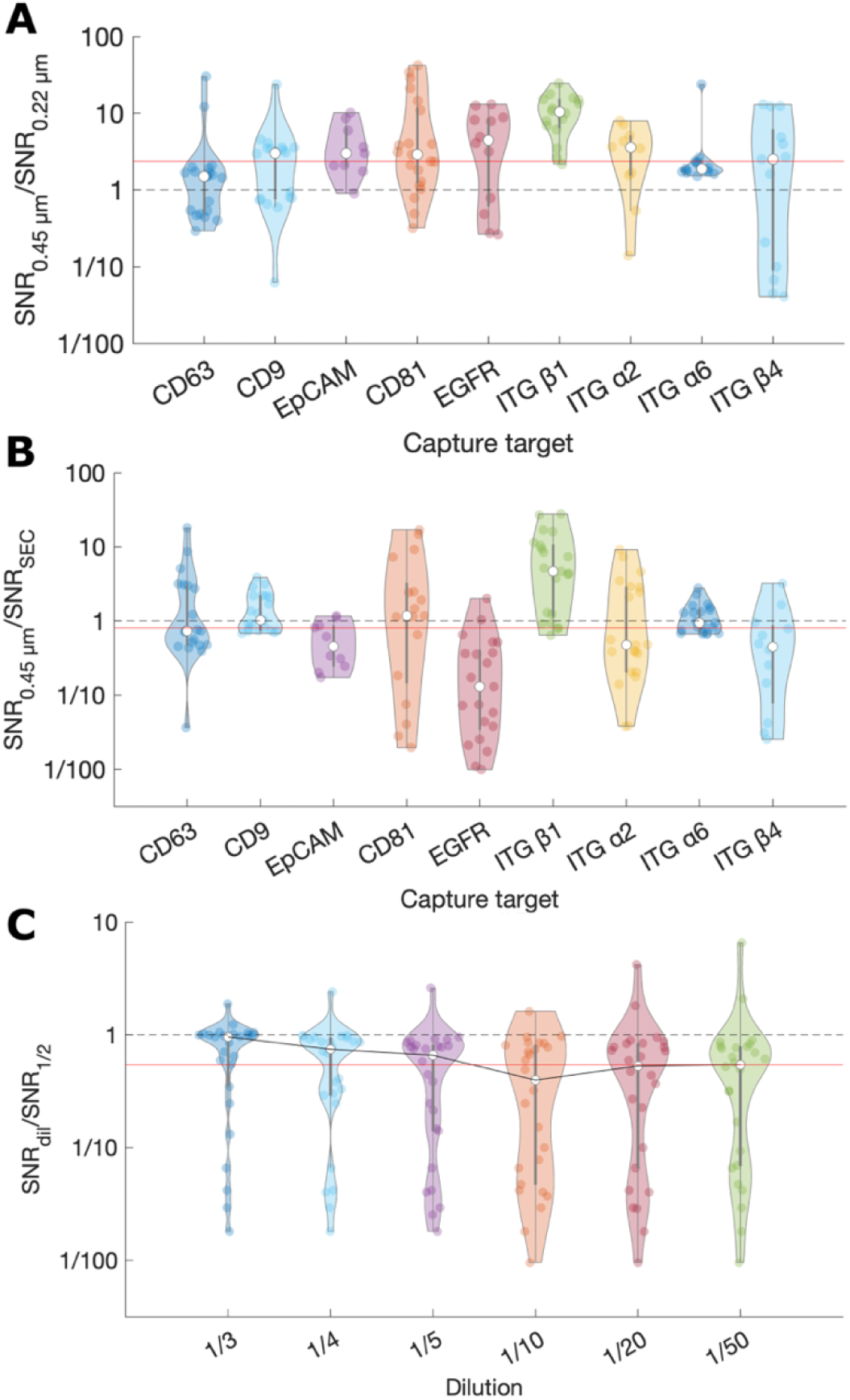
Optimization of a minimal plasma sample processing pipeline for EVPio. Comparison of SNRs obtained following (**A**) minimal processing using 0.22 or 0.45 μm filtration and (**B**) 0.45 μm filtration-based minimal processing and SEC purification of pooled plasma prior to EVPio analysis. (**C**) Drop in SNR with serial dilution for 0.45 μm filtration-based minimal processing. Plasma was minimally processed or SEC-purified and serially diluted, then EVs were captured based on select surface proteins and probed for their (**A**) CD63, CD9 and CD81 or (**B, C**) ITG β1, ITG β3 and CD9 content. Each dot represents the average of n = 10 technical replicates for a given (**A, B**) detection target and dilution or (**C**) co-expression pair. Red lines represent the average (**A, B**) or plateau (**C**) SNR ratio.

The working dilution factor applied upstream of EVPio analysis also impacts downstream SNRs. Greater EV heterogeneity and the presence of a complex biological matrix naturally result from the use of minimally purified plasma, leading to target concentrations spanning orders of magnitude and interactions at the EV surface (i.e., the EV corona^28,29^) impacting protein content and accessibility. While corona proteins can in themselves be informative, they can also be detrimental to affinity-based phenotyping efforts by shielding other potential biomarkers. Since dilution in PBS affects the binding equilibrium of corona proteins,^30^ we hypothesized that intermediate dilutions may allow one to compromise between both sources of informative proteins. EVPio assays performed on serially diluted, minimally processed pooled plasma EVs showed that detection SNRs are highest with a 1/2 plasma dilution in PBS but remain relatively stable at around 54 % of their maximal value past a 1/5 dilution, an effect that can be traced back to lower backgrounds compensating for decreasing signals at higher dilutions (**Figure 2C** and **Figure S3**). To avoid shifting the concentration of rarer targets outside of the detectable range in the final assay, we settled on a dilution range of 1/2-1/20 for further testing.

### Preliminary EVPio of CRCLM plasma EVs highlights inter-patient proteomic variability

The first step in developing a CRCLM-focused EVPio assay is to assemble a preliminary set of relevant proteins to target at the capture and detection steps. As an antibody microarray-based method, EVPio allows one to locally enrich EV subpopulations presumed to be informative and probe them *in situ*, enabling both combinatorial phenotyping and increased biomarker detection sensitivity.^23,31^ However, the success of this targeted phenotyping strategy hinges on the proteins included in the analysis recapitulating biomolecular variations relevant to the condition under study.^31^ We thus leaned on growing scientific understanding of how protein expression in CRC and CRCLM relates to disease progression, HGP presentation, liver function and both local and systemic immune reactions to build an EVPio panel likely to capture distinctive patterns between samples. To increase the odds of finding HGP-specific patterns, the initial target selection spans a wide array of candidate proteins, from more general EV and cancer-related proteins to promising HGP-related markers (**Table 1**).

**Table 1.** Protein targets evaluated for inclusion in the EVPio-CRCLM panel. Candidate proteins are spread across categories reflecting distinct aspects of the disease, from cell transformation to immunity and liver function.

| Category | Target | Capture/<br>Detection | Rationale |
| --- | --- | --- | --- |
| General EV | CD9 | Both | Common EV marker<sup>35</sup> |
|  | CD63 | Both | Common EV marker<sup>35</sup> |
|  | CD81 | Both | Common EV marker<sup>35</sup> |
|  | CD147 | Capture | Abundant in larger EVs (ectosomes) of cancer cell provenance<sup>36</sup>; pro-angiogenic properties<sup>37</sup> |
|  | LAMP1 | Capture | Late endosome/lysosome marker<sup>38,39</sup> |
|  | Syntenin-1 | Detection | Enriched in small EVs<sup>35,40</sup> |
|  | HSP70 | Detection | Commonly found in EVs, with increased abundance under stress<sup>41</sup> |
| Immune cells | CD11b | Capture | Regulator of myeloid cell function; promotes pro-inflammatory macrophage polarization and anti-tumor immunity<sup>42</sup> |
|  | CD68 | Capture | Over-expressed in tumor-associated macrophages and tumor cells; high expression correlates with higher tumor grade<sup>43</sup> |
|  | CD15 | Capture | Overexpression is related to lower T cell tumor infiltration, targeted therapy resistance and poorer outcomes in metastatic CRC<sup>44</sup> |
|  | CD19 | Detection | Marker of B cell activation; high expression correlates with better prognosis in CRCLM<sup>45</sup> |
|  | CD39 | Detection | CD39 upregulation contributes to tumor immune evasion and is related to poor outcomes in CRC and many other cancers<sup>46</sup> |
|  | S100A9 | Detection | Expressed by macrophages and neutrophils at the tumor interface in replacement CRCLM; showed potential in stratifying HGPs<sup>47</sup> |
|  | C5AR1 | Detection | Associated with tumor inflammation, recruitment of tumor-associated macrophages and tumor growth<sup>47,48</sup> |
|  | FERMT3 | Detection | Decreased expression in CRC; gain of expression found to enhance anti-tumor immunity and reduce chemoresistance<sup>49</sup> |
| Hepatocytes | HSA | Capture | Marker used to differentiate hepatocellular carcinoma from CRCLM<sup>50,51</sup>, but also found expressed in ~50% of cancers of the gastrointestinal tract<sup>52</sup> |
| Tumor cells | CLDN-2 | Both | Promotes CRC metastasis; marker of the replacement HGP<sup>32</sup> |
|  | EpCAM | Capture | Epithelial tumor marker; high expression in tissues and EVs related to poor outcomes<sup>53</sup> |
|  | CD82 | Both | Tumor metastasis suppressor; high expression correlates with better prognosis in CRC<sup>54</sup> |
| | ITG β1 | Both | Commonly secreted into EVs, with level increasing in cancer and with disease stage<sup>55</sup> |
|  | ADAM10 | Capture | Regulator of the NOTCH pathway; overexpression in CRC correlates with poor prognosis<sup>56</sup> |
| | CD44 | Capture | Involved in cell motility and often upregulated in cancer<sup>57</sup>; EV CD44 can interact with ITG $\alpha 6 \beta 4$ in liver cells to promote metastasis<sup>58</sup> |
| | ITG $\alpha 6$ | Both | Overexpression (in combination with ITG $\beta 4$) is associated with aggressiveness and poor outcomes<sup>59</sup> |
| | ITG $\beta 4$ | Both | Overexpression (in combination with ITG $\alpha 6$) is associated with aggressiveness and poor outcomes<sup>59</sup> |
| | ITG $\alpha v$ | Detection | EV expression of ITG $\alpha v \beta 5$ is associated with liver metastasis in tumor models<sup>60</sup> |
| | ITG $\beta 5$ | Capture | EV expression of ITG $\alpha v \beta 5$ is associated with liver metastasis in tumor models<sup>60</sup> |
| **Lipid processing** | ApoF | Detection | Regulator of cholesterol transport; EV-associated ApoF is a marker of colorectal neoplasia<sup>34</sup> |
|  | ApoA4 | Detection | Potential marker for HGP-based stratification<sup>47</sup> |
|  | CETP | Detection | Lipid transfer protein with increased activity in CRC patients<sup>33</sup> |

To confirm the suitability of the platform for the analysis of biobanked CRCLM patient plasma samples and trial the CRCLM-focused target panel, we carried out EVPio assays on a preliminary set of one sample from each rHGP and dHGP, as well as a liver cyst sample for comparison purposes (**Figure 3** and **Figure S4A-B**). The pairwise signal ratios between samples, presented in the bottom row of **Figure 3**, highlight how EVPio can detect inter-patient proteomic variations, including potential HGP-related patterns to be further investigated. For instance, FERMT3 has a stronger overall signal in both CRCLM samples relative to the cyst control but appears particularly highly expressed in the rHGP patient when combined with CLDN2, HSA and CD15. CLDN2, a tight junction protein associated to the rHGP,^32^ is predictably found to be more prevalent in the rHGP compared to the dHGP sample but interestingly even more abundant in the cyst control, especially when co-expressed with EpCAM. Tetraspanin- and EpCAM-paired CETP, a lipid transfer protein,^33^ as well as HSA- and CD9-associated ApoF, a regulator of cholesterol transport,^34^ show the strongest dHGP stratification potential of this introductory dataset. Overall expression patterns remain fairly consistent down to a 1/20 plasma dilution (**Figure S4C-D**), with SNRs settling at 42% of the median value obtained for ½ plasma (**Figure S4F**).

**Figure 3.**
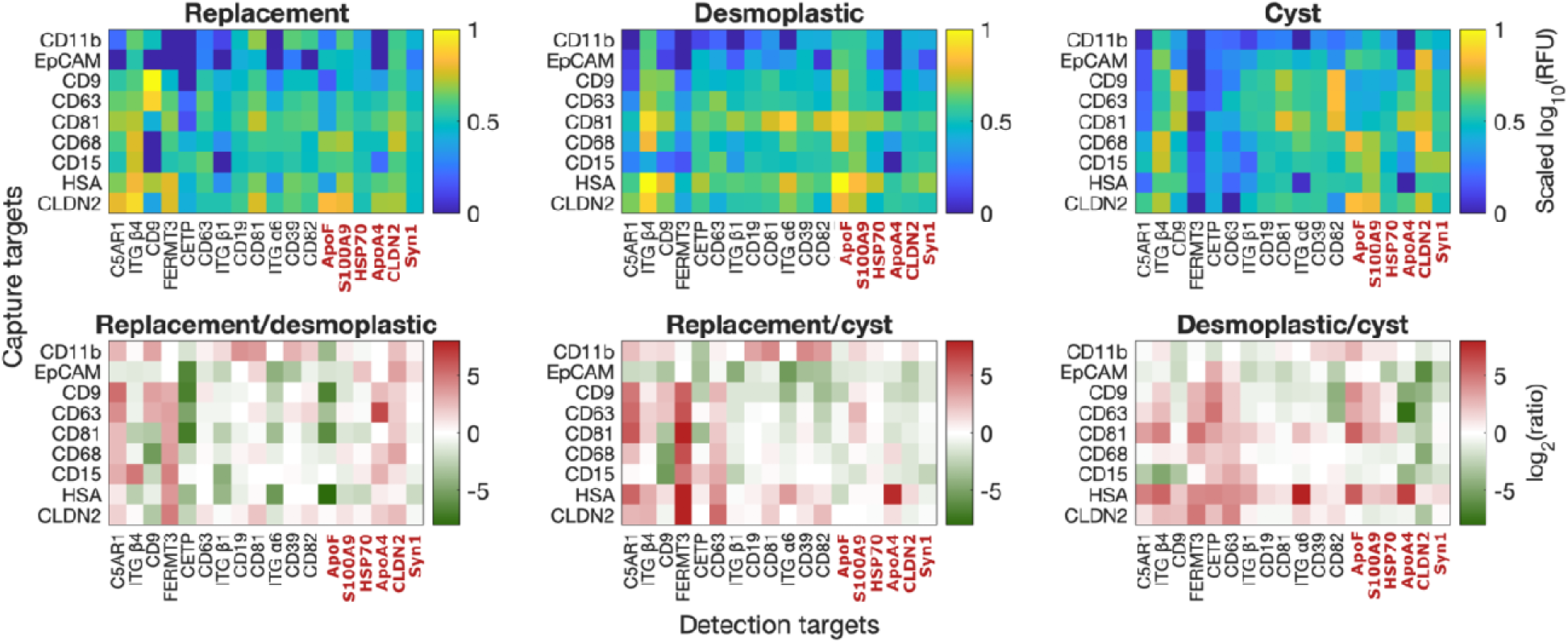
Preliminary phenotyping of patient plasma EVs highlights how EVPio can identify protein co-expression differences across frozen biobank samples without the need for an AR step. Co-expression (*upper* row) and ratio (*bottom row*) heatmaps of a set of *n* = 3 patient samples, one of each presentation (rHGP, dHGP and cyst), assayed at ½ dilution in PBS. Ratiometric analysis of this feasibility cohort provides early co-expression patterns for further exploration in larger cohorts, notably higher FERMT expression in rHGP, as well as CETP and ApoF expression in dHGP, especially when combined with tetraspanins.

A key feature of EVPio is the ability to probe both inner and outer EV proteins at the detection step. To evaluate whether the analysis of plasma EVs requires specific protocol adjustments, we assessed the effect of AR treatment on the signal of the six inner protein targets included in the panel (**Figure S4E**; inner targets are highlighted in red in **Figure 3**).

Unexpectedly, there was no significant improvement in the SNR of inner detection targets upon AR treatment (median 26% decrease in SNR), a trend at odds with previous observations in cell line EVs^23^. It is possible that biobanked plasma, being stored frozen for long periods and then thawed, contains EVs with reduced membrane integrity,^61^ enabling antibody-based detection of intravesicular antigens in the absence of a dedicated membrane-destabilizing step.

Another assay parameter that directly impacts assay signals and benefits from adjustment for plasma EV phenotyping is the capture antibody concentration used at the microarray printing step. The 100 μg/mL concentration used in the original EVPio assay was optimized for the capture and analysis of purified cell line EVs.^62^ Since plasma EV analysis requires the detection of low-abundance subpopulations in the presence of increased noise from minimal processing, we explored whether using a higher printing concentration of 200-300 μg/mL could substantially increase detection sensitivity. When using ratios to compare subpopulation signals (**Figure S5A**) and SNRs (**Figure S5B**) from EVPio assays run on 100 μg/mL and 200-300 μg/mL antibody microarrays, we found the later to yield overall stronger signals, with a 1.9× higher median signal. The impact on SNRs was less straightforward and more target-dependent, but the higher printing concentration was associated with pronounced SNR improvements (∼1.2-32×) for rarer, disease-specific subpopulations like CD68^+^, CD15^+^, CETP^+^ and EpCAM-S100A9^+^ EVs.

In light of these preliminary data, the final plasma EVPio workflow omits the AR step, uses antibody microarrays printed at ∼200-300 ug/mL, and requires samples diluted 1/2-1/20 in PBS, with the dilution factor chosen to balance signal intensity and sample availability.

### EVPio of a CRCLM pilot cohort guides refinement of an HGP-discriminating protein biomarker panel

We next combined the updated EVPio workflow and our preliminary target list to screen for the most differentially expressed proteins in a pilot cohort of nine (four rHGP, two dHGP and three cyst) patients (**Table S3**, **Figure S8**) as an intermediate pruning step upstream of a larger statistical learning-driven target selection. Exploratory biomarker studies commonly face challenges related to limited patient sample availability, which restricts the breadth of biomarker candidates that can be assessed.^31,63^ Preemptively removing extraneous panel targets based on insights from a pilot study narrows the feature-to-sample size gap, decreasing downstream dataset dimensionality and improving algorithm performance. This approach also mitigates the experimental challenges associated with running a large panel on dozens of samples in parallel, while lowering material and reagent costs.

We evaluated the suitability of biomarker candidates (**Table S1**) for inclusion in the final EVPio-CRCLM panel by assessing their 1) signal intensity and robustness (**Figure 4A-B**), 2) redundancy via pairwise correlation analysis (**Figure 4C-D**) and 3) HGP segregation potential through effect size estimation (**Figure 4E-F**). The large number of co-expression pairs (342) included in the pilot phenotyping (**Figure S6**), combined with the EVPio assay design only allowing single-antibody panel adjustments, limited the actionability of granular analysis at the antibody pair level (**Figure S6**, **Figure S7** and **Figure S9**). Consequently, we appraised putative target proteins for subpopulation capture (**Figure 4A, C, D**) and cargo probing (**Figure 4B, D, F**) independently by aggregating signals across detection and capture targets, respectively.

**Figure 4.**
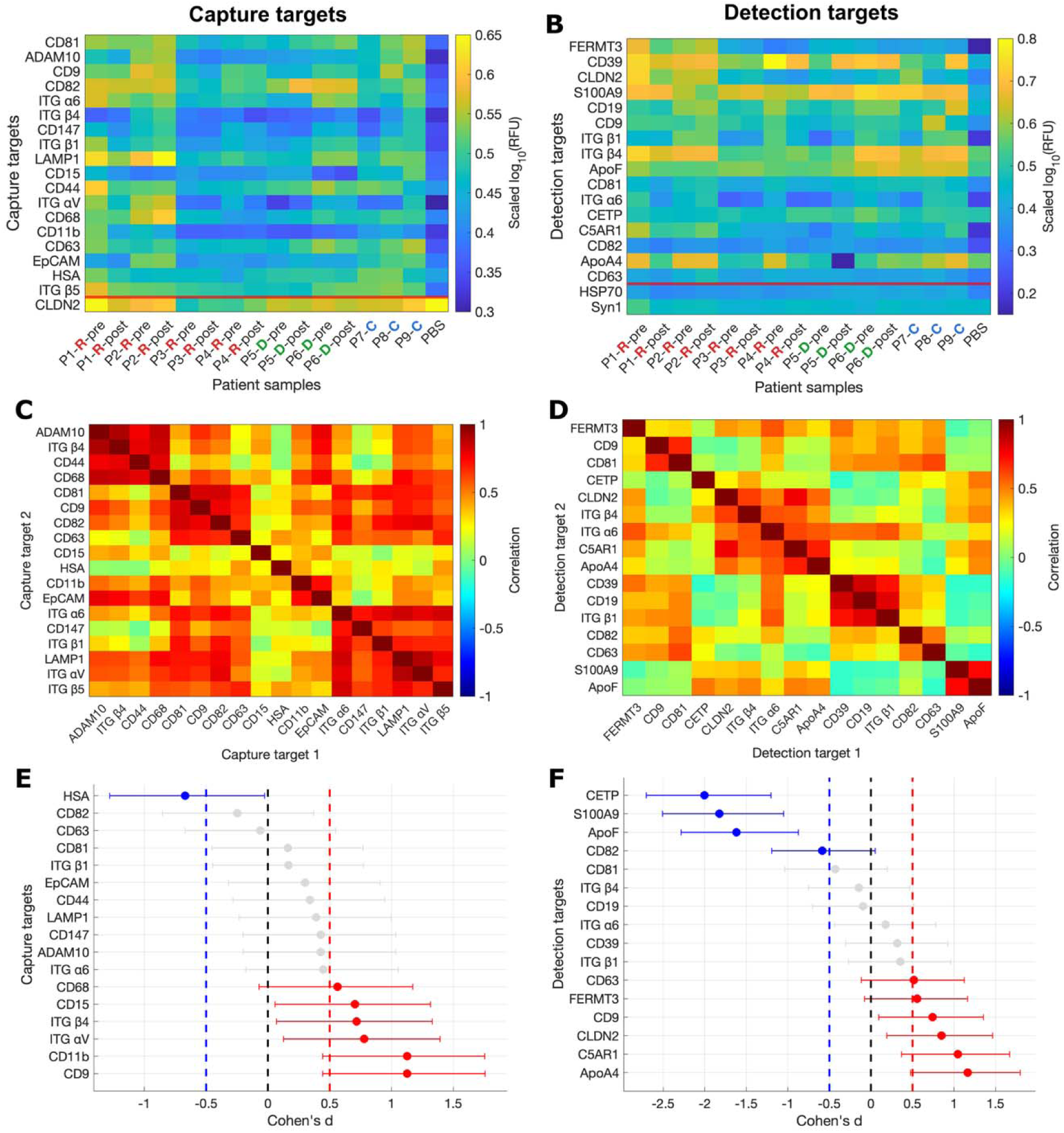
Selection of capture and detection targets for EVPio-CRCLM based on signal quality, correlation and effect size analysis of a pilot set of 15 samples from nine patients (P1-P9) across rHGP (**R**), dHGP (**D**) and cyst (**C**) presentations, including samples before (pre) and after (post) resection for rHGP and dHGP patients. Capture (**A**) and detection (**B**) signal, aggregated by taking the median over all marker-containing pairs and sorted by SNR, where the noise is the PBS negative control. The SNR threshold of 3 is represented as a red line. Clustered pairwise correlation map of (**C**) capture and (**D**) detection targets, calculated from the signal of all pairs containing the target. Capture targets (**C**) form thematic clusters dominated by proteins related to EV biogenesis (CD81, CD9, CD82, CD63), cancer and inflammation (ADAM10, ITG β4, CD44, CD68) as well as adhesion and integrins (ITG IZ6, CD147, ITG β1, LAMP1, ITG IZv, ITG β5), while detection targets (**D**) groupings reflect EV biogenesis and immunity (CD39, CD19, ITG β1, CD82, CD63, FERMT3, CD9, CD81) versus cancer and liver function (CLDN2, ITG β4, ITG IZ6, C5AR1, ApoA4, S100A9, ApoF). Cohen’s *d* effect size plots of (**E**) capture and (**F**) detection targets. Targets higher in rHGP and dHGP samples are represented in red and blue, respectively. Error bars show the spread of one standard deviation. A threshold of Cohen’s *d* = 0.5, pictured as a dashed line, was used to guide target selection. When considering the median over all pairs containing them, six capture targets (CD68, CD15, ITG β4, ITG IZv, CD11b, CD9) and six detection targets (CD63, FERMT3, CD9, CLDN2, C5AR1, ApoA4) were found to be more highly expressed in pilot rHGP patients, while one capture target (HSA) and four detection targets (CETP, S100A9, ApoF, CD82) had increased expression in dHGP subjects.

Panel targets were first sorted by average SNR and signals displayed as heatmaps, highlighting robust subpopulations showcasing endosomal (LAMP1^+^, CD9^+^, CD81^+^), cancer (CD44^+^, ITG α6^+^) and antitumor immunity (CD82^+^) markers (**Figure 4A**). Detection targets with strong signals and low noise included candidate HGP biomarkers S100A9 and CLDN2, immune evasion marker CD39 and lipid processing protein ApoF, among others (**Figure 4B**). Only one capture target and two detection targets fell below the SNR threshold of 3, leading to their exclusion from further analysis. Pairwise correlation coefficients (**Figure S7**) were next calculated and clustered for the capture (**Figure 4C**) and detection (**Figure 4D**) panels separately, emphasizing distinct groupings of associated proteins, including clusters enriched in tetraspanins, cancer markers and immunity-related proteins. Capture targets showed overall higher intercorrelation than detection targets, with CD15 and HSA standing out as having the weakest association with other subpopulations. The tetraspanin- and integrin-dominated groups demonstrated strong mutual correlation, while the tightest cluster (correlation coefficients ∼0.75-0.9) comprising CD44, ADAM10, CD68 and ITG β4—all related to tumor aggressiveness^43,56,59^—correlated more modestly to other subpopulations. Notable detection target associations include immunomodulatory proteins CD39,^64^ ITG β1^65^ and CD19^66^, and inflammation and immune infiltration markers CLDN2^32^ and C5AR1^48^. To identify targets with HGP separation potential, we calculated Cohen’s *d*, a standardized measure of effect size that takes into account variation around the group means, and can thus be calculated with a confidence interval.^67,68^ A threshold of *d* = 0.5, corresponding to a medium effect, highlighted 7 capture (**Figure 4E**) and 10 detection (**Figure 4F**) targets with good preliminary HGP stratification. Pilot patients with a rHGP had considerably more EVs bearing CD9, CD11b, ITG αV, ApoA4, C5AR1 and CLDN2, while dHGP patients had an overrepresentation of EVs expressing HSA, CETP, CD82, S100A9, and ApoF. While the bulk of these expression patterns agrees with previous findings where available,^32,47^ some are more surprising, such as the strong apparent relationship between CD9 and the rHGP—hinting at possible HGP-related alterations of the endosomal EV biogenesis pathway.

We selected the final EVPio-CRCLM panel by incorporating insights from all three analyses detailed above. Retained capture targets include CD15 and HSA, for their non-overlapping subpopulation contributions; LAMP1, CD63 and CD147 for their representation of various EV biogenesis pathways; and CD44, ITG α6, CD11b and EpCAM to cover cancer- and immunity-associated variation. For detection targets, CD9, CD39 and ITG β4 were selected for their robust signals and moderate interpatient variability; C5AR1, FERMT3, CETP, S100A9 and ApoF were chosen for their strong HGP stratification potential within this pilot dataset, as well as their emerging roles in CRC/CRCLM biology (**Table 1**).

### EVPio-CRCLM of a 58-patient cohort enables HGP identification with 75.9% cross-validated accuracy

The final iteration of EVPio-CRCLM (**Table S1**) was leveraged to assess 81 protein co-expression pairs on plasma EVs (**Figure S16**) from a cohort of 58 CRCLM patients, as well as nine cyst patients for comparative purposes (**Figure S11**). Unlike the pilot cohort, this final cohort includes many patients with liver metastases of mixed HGP composition, where the non-dominant HGP represents a non-negligeable proportion of the resected tissue (**Table S3**). Until recently, international consensus guidelines for HGP scoring set a 50% cutoff to assign a dominant HGP.^69^ However, the latest set of guidelines, published in 2022, builds on emerging prognostic insights to propose that the presence of any non-dHGP (rHGP or pHGP) should be the deciding scoring factor, essentially sorting CRCLM patients into dHGP and non-dHGP categories.^6^ Consistent with recent findings, we found that applying this Rotterdam cutoff^70^ (with rHGP designating the non-dHGP category) as opposed to the “proportional” 50% cutoff, led to noticeably better overall class discrimination performance of candidate co-expression pairs, with on average 4.8% larger standardized effect sizes (Cohen’s *d*) and 43% more features with |*d*| *>* 0.2 after noise-based thresholding (**Figure S12**).

Of the 81 candidate protein co-expression pairs (hereon also referred to as features/variables) measured, 40 had sufficient signal quality and robustness to be retained for classification (**Figure S11**). Feature selection based on class discrimination performance further refined this set to 8 candidate co-expression pairs as the foundation for classification (**Figure 5A**). Notably, protein marker CLDN2 and C5AR1 appear in multiple retained pairs, in line with known association with HGP-specific processes.^32,47^

**Figure 5.**
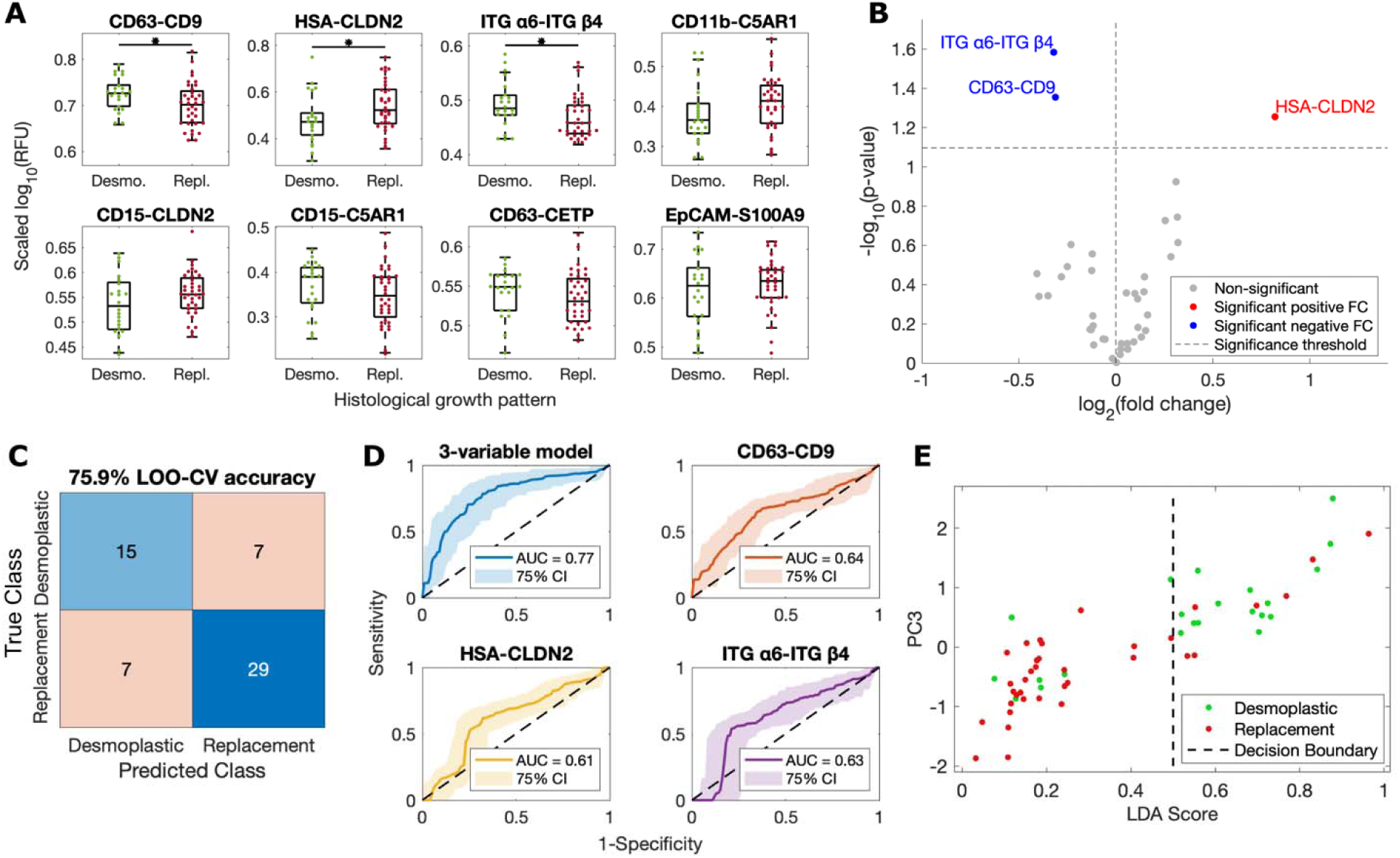
EVPio phenotyping and LDA-based HGP classification of the 58-cancer patient subset of the discovery cohort. (**A**) Dot plots of the most differential co-expression combinations based on ROC values. (**B**) Volcano plot of remaining protein pairs after noise-based thresholding. Three protein pairs have significantly (*p*-value < 0.08) higher expression in rHGP and dHGP samples and are shown in red and blue, respectively. (**C**) Confusion matrix of a three-variable LDA model optimized from the pruned variable set yielding a 75.9% cross-validated (leave-one-out, LOO) accuracy. (**D**) ROC curves of the selected three-variable LDA model compared to that of univariable models trained on each of the final model’s composite co-expression biomarker pairs. The ROC curves represent the median of 100 bootstraps with 75 percentile confidence intervals represented as shaded areas. (**E**) 2D representation of the three-variable LDA model decision boundary, where each sample is plotted according to its LDA score and third principal component (PC3).

To develop a robust classification model, we applied linear discriminant analysis (LDA) with recursive feature elimination (RFE) (see Methods), supplemented by significance and effect size assessment (**Figure 5B** and **Figure S12**). Given the considerable feature intercorrelation (**Figure 5C-D**) and the *n* = 58 cohort size, a conservative feature set size of *d* = 3 was chosen to mitigate overfitting risks.^71^ The optimal model, incorporating CD63-CD9, HSA-CLDN2 and ITG α6-ITG β4, achieves 75.9% accuracy under leave-one-out (LOO) cross-validation (**Figure 5C**). Compared to analogous LDA models trained on individual composite features, the optimal 3-feature model demonstrates superior cross-validated performance, quantified through higher accuracy and consistency across classification thresholds (AUC = 0.77 > 0.61-0.64, **Figure 5D**).

Plotting LDA scores from this optimal model against the third principal component of the composite features reveals clear class separation, with over two-thirds of the rHGP patients clustering in a narrow band around LDA score ∼ 0.2, while dHGP patients span a broader range of more moderate values (0.5-0.8) (**Figure 5E**). Despite this overall separation, some samples cluster closely with the opposite class. Principal component analysis (PCA) and clustering of the noise-filtered dataset reveals substantial similarity between the two classes, visualized as intertwined clusters (**Figure S13**). This overlap, along with the uncertainty underlying 75.6% classification accuracy, likely reflect high inter-patient variability and shared biological mechanisms between subtypes, which may contribute to classification challenges. Nevertheless, our 3-feature LDA model achieves markedly improved separation compared to unsupervised clustering, with considerably less information. Altogether, our model demonstrates promising separation of CRCLM patients based on liver metastasis HGP and identifies key biomarkers and protein interactions with potential for non-invasive HGP typing.

## Discussion

Here, we describe the development of a targeted EVPio-CRCLM assay and analytic framework for HGP-based classification of CRCLM patients, aimed at enhancing disease assessment and guiding treatment selection. Building upon our established EVPio antibody microarray-based workflow, we added a high-throughput sample preparation pipeline that minimizes sample loss and can easily be performed in parallel with other preparatory steps like microarray printing. We then leveraged the multiplexed nature and wide antibody compatibility of EVPio, which is particularly well suited to the design of bespoke, condition-specific assays, to iteratively construct a CRCLM-focused biomarker panel. In a cohort of 58 CRCLM subjects, we measured 81 protein co-expression pairs—divided into nine immunocaptured subpopulations on which nine additional markers are probed—with as little as 30 μL of plasma per patient. An LDA-based classifier trained on the resulting phenotyping data achieves 75.9% accuracy in sorting patients based on HGP by combining measurements from just three protein co-expression pairs.

To process plasma samples for EVPio analysis, we wanted a straightforward, quick, scalable workflow that strikes a balance between EV retention (signal) and impurity (noise) removal, which we quantified using the SNR. While more complex techniques like SEC and ultracentrifugation yield higher-purity samples, EVPio features like optimized surface passivation, orthogonal barcode-enabled low cross-reactivity and multiplexed detection, spatially defined detection areas (antibody microspots) and reliance on dual antibody recognition allowed us to circumvent time-consuming pre-processing and lower sample volume requirements, with comparable downstream SNR. Other than signal quality, another concern related to sample processing is the introduction of biases in subsequent measurements, such as an altered EV size distribution (for methods based on physical properties) or an overrepresentation of specific phenotypes (for affinity-based methods). While SEC or filtration avoids direct phenotypic bias, protein profiles can be indirectly affected as a result of correlations between protein abundance and EV size (e.g., due to biogenesis differences^35^). Comparison of our minimal processing pipeline to our previous SEC-based protocol revealed noticeable SNR variations in two of nine subpopulations (**Figure 2B**), suggesting that expression of the corresponding protein markers might be confounded with sample properties impacted by pre-processing, e.g., EV size distributions. Notably, their respective SNRs diverge in opposite directions, underscoring that both processing methods have the potential to introduce biases, which warrants careful consideration in experimental design and data interpretation.

For an affinity-based platform like EVPio, insights derived from EV phenotyping depend critically on the judicious selection of targets. While existing literature (see **Table 1**) provided a foundation for constructing a preliminary panel, data on target co-expression patterns and their relationship to CRCLM HGPs remain sparse. To address this, we iteratively refined our panel by excluding antibodies with poor specificity or targets lacking differential signals across HGPs, while retaining those with promising associations. A key limitation of this approach lies in the biological complexity of CRCLM: pairs sharing a common capture or detection target exhibited wide expression variability and even opposing HGP associations, making it more challenging to curate the target panel at the single protein level. For example, in our pilot study, CD15-ITG β4 correlated with rHGPs, whereas HSA-ITG β4 was enriched in dHGP samples (**Figure S9**). Furthermore, insights from early screening were constrained by limited sample availability—only nine subjects could be included in the pilot, and all were chemotherapy-naïve cases with a dominant HGP, potentially biasing preliminary findings towards an idealized subset unrepresentative of real-world heterogeneity.

The preliminary and pilot studies were also critical to the refinement of the plasma-compatible updated EVPio protocol, offering opportunities to tune assay parameters such as the AR step, sample dilution factor and microarray printing specifications. While AR proved essential for detecting intravesicular proteins in EVPio analysis of fresh SEC-purified cell line EVs,^23^ we observed no clear benefit in its application to frozen biobanked plasma samples (**Figure S4E**). We speculate that this discrepancy arises from differences in EV integrity and density: frozen storage is known to compromise lipid bilayers,^61,72^ and the comparative rarity of target EVs in plasma due to increased heterogeneity may reduce local crowding and antigen density at microarray spots, together potentially easing antibody access and binding.^73,74^ AR may thus yield diminishing returns as such low-abundance conditions compound the vulnerability of fragile EV samples to heat-induced loss.^23^ Consequently, we did not use AR in the final EVPio-CRCLM workflow. In parallel, we optimized the sample dilution factor to balance the competing priorities of signal quality, dynamic range, and sample conservation. Higher dilutions (e.g., 1/10) improved signal-to-noise ratios (SNRs) for select targets, potentially by reducing nonspecific background noise or improving antibody accessibility to sterically hindered epitopes (e.g., corona-shielded proteins, which can be exposed through sample dilution or buffer exchange^30^). However, most co-expression pairs exhibited ≥ 50% SNR reduction at 1/10-1/20 dilution, with some falling even more sharply (**Figure S4F**). This decline likely reflects diminishing target abundance below detection thresholds, particularly markers present in EVs with low copy numbers. To minimize sample and reagent consumption while retaining detectable signals across most targets, we selected an intermediate 1/5 dilution, prioritizing broad applicability over optimizing for individual co-expression pairs.

The final EVPio-CRCLM study probed the best candidate markers from the previous two preparatory studies in a larger cohort of 58 CRCLM patients (**Figure S11**). We sought to identify the optimal combination of markers to stratify patients by HGP. Given the exploratory nature of the current study, we prioritized hypothesis generation by selecting a slightly relaxed significance threshold (*p* < 0.08) for feature testing to avoid missing relevant biological signals due to limited statistical power. We paired this with a robust multi-step feature selection pipeline (SNR/CV-based noise-based filtering, outlier removal, ROC-based feature ranking and cross-validated RFE; see Methods) to emphasize pairs with high quality signal, low intercorrelation and high individual HGP stratifying potential, bolstering model robustness and mitigating overfitting risk. The top-performing LDA classifier (75.9% accuracy) integrated three co-expression pairs: CD63-CD9, HSA-CLDN2, and ITG α6-ITG β4. These pairs span general EV, hepatocyte-like and tumor-stromal markers (**Table 1**), likely reflecting distinct biological pathways underlying HGP divergence. The elevated HSA-CLDN2 in rHGP samples aligns with our prior work identifying CLDN2—a mediator of tumor-hepatocyte adhesion and liver colonization—as a hallmark of rHGP, which was also found to be uniquely expressed in EVs from rHGP plasma samples using ultracentrifugation followed by western blotting.^32^ The co-expression of HSA, a hepatocyte-specific antigen,^50,51^ may reflect the structural mimicry of liver parenchyma by rHGP tumors,^6,7^ enabling immune evasion and integration into hepatic tissue.

Together, HSA-CLDN2 thus possibly captures the dual adhesive and stealth mechanisms driving rHGP pathology. The predictive power of CD63-CD9, while initially unexpected, might be a result of stromal crosstalk in dHGP lesions. Activated immune cells and cancer-associated fibroblasts (CAFs) in dHGP stroma are prolific sources of tetraspanin-bearing EVs,^75,76^ which mediate cell-cell communication and extracellular matrix remodeling. Similarly, the contribution of EV ITG α6β4—a laminin-binding integrin linked to invasion and stromal remodeling^59^—to dHGP identification could be related to its role in CAF-driven fibrosis and metastatic niche formation.^60^ Hence, through stringent predictor selection and by potentially leveraging HGP-specific biology—rHGP liver mimicry (HSA-CLDN2) and dHGP stromal activity (CD63-CD9, ITG α6β4)—this three-part signature achieves preliminary classification with encouraging accuracy (AUC = 0.77) despite inherent cohort limitations.

Inter-patient variability within each HGP class poses a significant challenge to accurate classification, particularly in a moderately-sized cohort like ours (*n* = 58). Comparison of LDA and PCA projections (**Figure 5E** and **Figure S13**) reveals that stratifying variance, primarily captured in the third principal component, is overshadowed by unrelated sources of variability dominating the first two components. Despite this subtler separation signal, LDA, as a supervised learning method, can seek HGP-specific patterns to amplify class distinctions while accounting for variability within each group. However, classification performance varied across classes, with dHGP samples showing a higher misclassification rate (32%) compared to rHGP samples (19%) (**Figure 5C**). This disparity likely reflects both biological and methodological factors: the class imbalance (36 rHGP vs. 22 dHGP) derived from Rotterdam HGP grading (pure dHGP vs. any proportion of non-dHGP), which mirrors real-world clinical populations where pure dHGPs are uncommon,^32,70^ may limit the model’s ability to resolve heterogeneity within the smaller dHGP subgroup. At the onset of this study, the Rotterdam cutoff had not yet reached consensus, limiting our ability to optimize sample inclusion criteria to ensure balanced classes and equal training opportunity for dHGP and non-dHGP samples. This challenge is compounded by the higher variability of putative dHGP-associated pairs (CD63-CD9, ITG α6β4), as seen in their proportionally lower fold changes (∼1.25×) compared to that of tentative rHGP marker pair HSA-CLDN2 (1.77×). Echoing this dichotomy, rHGP specimens cluster tightly on the LDA axis (**Figure 5E**), while dHGP samples exhibit greater dispersion. These results could stem from robust HSA-CLDN2 expression in non-dHGPs even with minor rHGP foci, consistent with the prevailing hypothesis of rHGP dominating CRCLM biology.^70^ Similarly, increased but variable expression of CD63-CD9 and ITG α6β4 in the dHGP class might be reflective of the enhanced immune activation and stromal complexity central to the current dHGP paradigm.^6,77^

Closer examination of samples misclassified by our LDA classifier (**Figure S14**) reveals distinct biomarker patterns: dHGP samples misclassified as rHGP tend to exhibit elevated HSA-CLDN2 and reduced CD63-CD9, while misclassified rHGP samples show the inverse trend (higher CD63-CD9, lower HSA-CLDN2). Several factors may contribute to these discrepancies, including the universal use of neoadjuvant chemotherapy in our cohort. Variability in treatment response could alter EV profiles, particularly for markers linked to stromal activity. For example, chemotherapy-responsive rHGP lesions may display increased necrosis and immune-mediated fibrosis,^70^ elevating stromal EV signals (e.g., CD63-CD9) and increasing the likelihood of misclassification as dHGP. Conversely, misclassified dHGP samples with elevated HSA-CLDN2 could reflect spatial or temporal sampling biases. Since candidate rHGP-associated signals (i.e., HSA-CLDN2) seem to arise with even minimal rHGP components, tumors classified as 100% dHGP based on histopathological analysis might harbor hidden rHGP biology—possibly within new micrometastases or missed during sampling. This hypothesis aligns with preliminary findings that new metastases arising during or after preoperative treatment tend to display a rHGP phenotype,^8^ contrasting with the observed pattern of increased dHGP frequency within resected tumors post-chemotherapy.^78^ In this context, dHGP cases misclassified as rHGP could represent a high-risk subgroup with an increased likelihood of developing rHGP lesions and associated poor outcomes. While this is consistent with the prognostic rationale of the Rotterdam HGP grading system, the potential link between EV proteomic signatures and disease progression warrants further investigation in larger, longitudinal studies.

Another complexity in HGP-based modeling is the handling of mixed lesions, which hinges on the framework used to assign class membership. We compared the two predominant HGP grading systems: the proportional (50%) cutoff^69^ and the Rotterdam criteria^6^ (**Figure S15**). The Rotterdam framework improved the predictive power of putative stromal markers (CD63-CD9, ITG α6β4) and enabled the emergence of HSA-CLDN2 as a candidate rHGP-specific biomarker. This could stem from the Rotterdam criteria’s exclusion of specimens with even minimal rHGP components from the dHGP subtype, thereby reducing HSA-CLDN2 expression in the resulting (pure) dHGP class and minimizing within-class variability. In contrast, the proportional cutoff’s inclusion of mixed lesions in the dHGP class may dilute dHGP-associated biomarker signals (e.g., CD63-CD9, ITG α6β4) by boosting heterogeneity in immune-stromal phenotypes.

This study presents a preliminary liquid biopsy framework based on EV protein co-expression analysis for HGP classification in CRCLM. By integrating a high-throughput EVPio-CRCLM assay with supervised learning, we identified three co-expression pairs that collectively stratify HGPs with promising accuracy, highlighting the potential of combinatorial EV proteomics to recapitulate key features of the rHGP-dHGP dichotomy within liver metastases. While limited by modest sample sizes and homogeneous, idealized early cohorts, these findings demonstrate that EV phenotyping can provide insight into tumor-stroma dynamics beyond conventional histology, opening a window into non-invasive HGP typing. Serial sampling could further extend this application to longitudinal monitoring, facilitating real-time assessment of therapeutic response and early detection of disease recurrence.

Follow-up studies in larger cohorts will be essential to validate and strengthen the proposed biomarker signature, refining stratification accuracy and capture of CRCLM biology by disentangling confounders (e.g., treatment regimens^78^) and integrating complementary HGP-specific protein biomarkers (e.g., related to encapsulation and fibrosis in the dHGP^79^). Ultimately, EV-based scoring has the potential to transform HGP stratification from a static, post-resection histology metric into a minimally invasive preoperative assessment, guiding the selective use of anti-angiogenic therapies to maximize benefit while sparing non-responders from unnecessary pharmacological burden. More broadly, this step towards liquid biopsy-guided HGP classification lays the foundation for advancing precision oncology in CRCLM.

## Materials & Methods

### Cell culture

HT29 cells (ATCC® HTB-38) were cultured in Dulbecco’s Modified Eagle Medium containing 4.5 g/L D-glucose, L-glutamine, 110 mg/L sodium pyruvate (DMEM, Gibco), 10% fetal bovine serum (FBS, Gibco) and 1% penicillin-streptomycin (PS, Thermo Fisher Scientific), under constant humidity conditions at 37°C and with 5% CO_2_.

### Size-exclusion chromatography (SEC) purification of cell line and plasma EVs

#### Cell line EVs

HT29 cells were seeded into 3- or 5-layer TC-treated multi flasks (Falcon) and expanded for 24h before the culture media was refreshed with new media containing 5% exosome-depleted FBS (Gibco) and 1% PS, after what the cells were grown to 80% confluency without further media change. The media was then transferred to 50 mL tubes, centrifuged for 15 min at 400 g to remove large cell debris, syringe-filtered (pore size 0.22 μm, 33 mm diameter, MilliporeSigma) and subjected to successive 25-min spins at 4000 rpm in Amicon Ultra-15 centrifugal filters (100 kDa cutoff; Millipore Sigma) until concentrated to approximately 500 μL per 30 mL. This cell supernatant concentrate was applied onto qEV original columns (70 nm; Izon Science), followed by flushing with filtered (0.22 μm filter, MilliporeSigma) phosphate buffer saline (PBS, Gibco) and collection of the eluate in 500 uL fractions. Fractions 8 and 9 were combined and used as purified HT29 EV samples in subsequent experiments.

#### Plasma EVs

Pooled human plasma in sodium citrate (BioChemed) was centrifuged at 10,000 *g* for 10 minutes, then the supernatant was syringe-filtered (0.22 μm pore size, 13 mm diameter, MilliporeSigma). SEC purification was performed by applying 150 uL plasma per qEVsingle Gen 2 column (70 nm, Izon Science), flushing with filtered PBS, and collecting 170 uL fractions after discarding the 870 uL void volume. The first four fractions were combined for use as purified pooled plasma EV samples in subsequent experiments.

### Assessment of SEC-related EVPio SNR losses

HT29 EVs were SEC-purified and dyed with CFDA-SE (V12883, Invitrogen) at 1/250 concentration for 2h at 37°C, then desalted in 100 kDa Amicon centrifugal filters (MilliporeSigma) to remove excess dye before being diluted back to their original volume in PBS. The dyed HT29 EVs were then mixed 1:1 by volume with human pooled plasma or filtered PBS. SEC purification was performed as detailed above for pooled plasma, and the collected eluate concentrated down to 100 uL using 100 kDa Amicon centrifugal filters. Inkjet-spotted antibody microarrays targeting CD63, CD9, EpCAM, CD81, ITG α6 and ITG β4 (**Table S1**) were prepared as previously described,^23^ then used to capture EVs (incubation of 2h at room temperature [RT], followed by overnight at 4°C), which were then detected based on CFDA-SE fluorescence.

### Minimal plasma sample processing

#### Optimization

Pooled plasma was centrifuged at 10,000 *g* for 10 minutes, then the supernatant was syringe-filtered with either a 0.22 μm or 0.44 μm centrifuge filter (13 mm diameter, MilliporeSigma). A portion of each filtered plasma condition was then concentrated down ∼3.3x by volume using 100 kDa Amicon centrifugal filters. Each sample condition was serially diluted (1/2, 1/3, 1/4, 1/5, 1/10, 1/20, 1/50) in filtered PBS before being applied to inkjet-spotted antibody microarrays targeting CD63, CD9, EpCAM, CD81, EGFR, ITG β1, ITG α2, ITG α6 and ITG β4 and detected in triplex configuration using conjugated antibodies against CD63, CD81 and CD9 (**Table S1**) and complementary fluorescent branched DNA probes (in order of dye: Alexa Fluor 647, Cy3 and Alexa Fluor 488) following the published EVPio workflow^23^ without AR.

#### Comparison to SEC

Pooled plasma was centrifuged at 10,000 *g* for 10 minutes, then the supernatant was syringe-filtered with a 0.44 μm centrifuge filter (hereafter referred to as “minimal processing”). Half of the sample was SEC-purified as described above, then concentrated down to 150 μL using 100 kDa Amicon centrifugal filters. Each sample condition was serially diluted (1/2, 1/3, 1/4, 1/5, 1/10, 1/20, 1/50) in filtered PBS before being applied to inkjet-spotted antibody microarrays targeting CD63, CD9, EpCAM, CD81, EGFR, ITG β1, ITG α2, ITG α6 and ITG β4 and detected in triplex configuration using conjugated antibodies against CD9, ITG β1 and ITG β3 (**Table S1**) and complementary fluorescent branched DNA probes (in order of dye: Alexa Fluor 647, Cy3 and 488) following the published EVPio workflow^23^ without AR.

### EV sample characterization

The protein content (in mg/mL) of SEC-purified HT29 EVs was estimated using a NanoDrop 1000 spectrometer (ThermoFisher Scientific) in A280 mode. The concentration (particle/mL) of SEC- and minimally purified samples were assessed by Microfluidic Resistive Pulse Sensing (MRPS) (nCS1 particle analyzer, Spectradyne) or Nanoparticle Tracking Analysis (NTA) (NanoSight NS300, Malvern Panalytical).

### Antibody barcoding and branched DNA amplification

Antibody conjugation and fluorescent branched DNA probe assembly were carried out as previously described, using the 2-branch design.^23^ A list of DNA barcodes and corresponding antibody targets used in this study can be found in **Table S2**.

### Patient plasma sample collection and HGP typing

This study was conducted in accordance with the McGill University Health Centre (MUHC) Institutional Review Board (IRB) guidelines. Samples were collected through the MUHC Liver Disease Biobank, and prior written informed consent was obtained from all patients included in the study (protocol: SDR-11-066). Inclusion criteria: confirmed CRCLM disease categorized as dominantly dHGP or rHGP (for CRCLM patients) or confirmed presence of benign liver cyst(s) (for non-CRCLM group). Clinical data (summarized in **Table S3**) includes demographics and disease characteristics and was obtained from medical and hospital records.

Patient HGP proportions were determined by pathologists through the examination of hematoxylin and eosin (H&E)-stained sections, following consensus histopathology scoring guidelines.^69^ A total of 64 CRCLM patients, as well as 10 liver cyst patients, met the criteria and had adequate plasma sample volume for inclusion in this work (**Table S3**). CRCLM patients included in the preliminary and pilot assays were chemonaïve at the time of blood collection. CRCLM patients included in the main EVPio-CRCLM study received neoadjuvant chemotherapy treatment, which was discontinued five to six weeks prior to blood collection at the time of tumor resection. Plasma isolation was done by centrifuging blood samples twice at 2500 *g* for 15 minutes at RT.

### Preliminary EVPio-CRCLM assay development

#### Panel testing and assessment of AR effect

Minimally processed plasma samples from one rHGP, one dHGP and one liver cyst (non-cancer) patients (**Table S3**) were serially diluted (1/2, 1/10 and 1/20) in PBS, incubated on inkjet-spotted antibody microarrays targeting CD11b, EpCAM, CD9, CD63, CD81, CD68, CD15, HSA and claudin-2 and detected with barcoded antibodies and complementary fluorescent branched DNA probes (in order of probe dye: Alexa Fluor 647, Cy3, Alexa Fluor 488) in the following triplex configuration: C5AR1, ITG β4, CD9; FERMT3, CETP, CD63; ITG β1, CD19, CD81; ITG α6, CD39, CD82; ApoF, S100A9, HSP70; ApoA4, claudin-2, and sytenin-1 (**Table S1**, **Table S2**). The assay followed the EVPio workflow^23^ without AR, with the exception of inner targets ApoF, S100A9, HSP70, ApoA4, claudin-2 and syntenin-1, which were assayed with and without AR.

#### Impact of increased capture antibody spotting concentration

Minimally processed plasma samples from the same three patients were serially diluted (1/5 and 1/10) and incubated on inkjet-spotted antibody microarrays targeting the same proteins as in the previous experiment, but at both 100 and 200-300 μg/mL (depending on stock antibody concentration). Conjugated antibodies against CETP, S100A9, CD9 and CD39 (overnight incubation at 4°C following 2h incubation at RT) and complementary Cy3 or ATTO 565-bearing branched DNA probes were used for singleplex detection.

### Pilot EVPio-CRCLM

Minimally processed plasma samples from four rHGP patients, two dHGP patients and 3 liver cyst patients (**Table S3**) were serially diluted (1/5 and 1/10) in PBS, incubated on antibody microarrays (200-300 μg/mL print) targeting CD11b, EpCAM, CD9, CD63, CD81, CD68, CD15, HSA, claudin-2, CD147, LAMP1, ADAM10, CD44, CD82, ITG α6, ITG β1, ITG β4, ITG αV and ITG β5 (**Table S1**), then detected with barcoded antibodies (2h incubation at RT followed by overnight at 4°C) and complementary fluorescent branched DNA probes in the same triplex configuration as the preliminary assay testing. Five technical repeats (antibody spots) were assigned per condition. The assay followed the EVPio workflow^23^ without AR.

### Main (discovery) EVPio-CRCLM study

Minimally processed plasma samples from 36 rHGP patients, 22 dHGP patients and nine liver cyst patients (**Table S3**) were diluted 1/5 in PBS, incubated on antibody microarrays (200-300 μg/mL print) targeting CD15, HSA, LAMP1, CD63, CD147, CD44, ITG α6, CD11b and EpCAM, then detected with barcoded antibodies (2h incubation at RT followed by overnight at 4°C) and complementary fluorescent branched DNA probes (in order of probe dye: Alexa Fluor 647, Cy3, Alexa Fluor 488) in the following triplex configuration: FERMT3, CETP, CD9; C5AR1, CD39, ITG β4; ApoF, claudin-2, S100A9 (**Table S1**, **Table S2**). Ten antibody spots were assigned per condition, and the assay was repeated twice, for a total of 20 technical repeats, with the exception of S100A9 and CLDN2, which have 10 replicates each due to antibody batch performance consistency issues. The order of patient samples on the microarrays was newly randomized before each of the two assay repeats. The assay followed the EVPio workflow^23^ without AR.

### Data analysis

#### General data preprocessing

Data was extracted from the microarray images using Array-Pro Analyzer software (MediaCybernetics). Further processing, analysis and plotting were carried out using custom MATLAB (MathWorks) scripts written in-house, with the exception of violin plots.^80^ Briefly, raw fluorescence signal values were corrected for inter-channel bleedthrough using calibration data, then the mean raw fluorescent unit (RFU) value of each spot was background-corrected by subtracting the mean RFU value of its local background. All further calculations were performed on log-transformed data. Outliers within the technical repeats of each condition were identified and removed based on the median absolute deviation (MAD) method (outliers are defined as further than 3*MAD from the median). Measures of central tendency (mean, median) and dispersion (standard deviation [SD], MAD, percentiles) for each condition were calculated using the log-transformed values of its technical repeats. SNR values used for optimization were calculated by subtracting the mean value of the matched no-EV (PBS) control and dividing the result by the no-EV control SD. Bootstrapping of technical replicates was used to calculate the associated 95% confidence interval. For heatmap visualization, aggregated values were scaled based on the maximum log_10_(RFU) value of the corresponding fluorescence channel.

#### Pilot dataset: biomarker performance assessment

For signal intensity assessment, the mean capture target SNR was calculated by taking the median across all detections, and vice versa. Capture and detection targets were sorted by SNR (median of detectable occurrences, defined as having SNR > 1, i.e., at least one standard deviation above noise), and a cutoff of SNR = 3 was used assess signal robustness. Correlation analysis was then carried out on the retained targets. Correlation matrices were calculated separately for capture and detection targets using the entire set of signal values (every combo of every patient) for each capture or detection target, then clustered by the complete linkage hierarchical clustering method with a threshold of half the maximal linkage distance. Inter-class differences were evaluated using volcano (for expression pairs) and forest (Cohen’s *d*) plots (for individual targets). For the volcano plot, significance, quantified as the *p*-value, was derived from two-sample *t*-tests run on log-transformed RFU values, while the fold change was computed based on the geometric mean of each class. For the forest plots, calculations were based on the median in log space across all pairs containing a given target. Cohen’s *d* values were obtained as previously described^68^; the standard deviation interval was calculated based on formulas for the scaled non-central *t* distribution.^81^ Targets were then sorted by their *d* value, and an absolute threshold of *d* = 0.5 was used to guide target selection.

#### Main dataset: statistical analysis and HGP classification

The correlation between the two separate assay run datasets (using the calculated average raw RFU value in log scale for each condition) was assessed to ensure reproducibility (**Figure S10**). Datasets were then batch-corrected and combined using a MATLAB implementation of the ComBat data harmonization method.^82,83^ The resulting dataset includes *n* = 58 CRCLM patients and *p* = 81 features. Low-quality features (protein co-expression pairs), defined as having either a >30% median coefficient of variation across repeats or a SNR < 1.75 (median of detectable occurrences, defined as having SNR > 1, i.e., at least one standard deviation above noise) were filtered out of the dataset. Outlier removal (based on the interquartile range [IQR] method: outliers are more than 1.5*IQR from the lower/upper quartile) was next performed on the resulting combined dataset on a feature-by-feature basis (i.e., across patients). Missing values were replaced using kNN imputation with *k* = 3 for downstream analytical compatibility. Cohen’s *d* values were calculated for each protein co-expression across the rHGP and dHGP groups. A threshold of *d* = 0.4 was applied to highlight promising features.

The filtered dataset of *n* = 58 CRCLM patients and *p*′ = 40 features (**Figure S11**) was z-score normalized and used to develop a binary classification model. A pre-selection of *p*″ = 8 features was performed based on ranking of their receiving receiver operating characteristic (ROC) performance (Z-values), with a correlation weight of 0.5 (which limits the intercorrelation of features included in the set). Recursive feature elimination, using 1−the area under the ROC curve (AUC) as minimization criterium, was then used conjointly with linear discriminant analysis (LDA) and *k-*fold cross-validation (*k* = 5) to build a model using only *p* = 3 predictors. The hyperparameters (γ, δ) of the model were then optimized (MATLAB’s automatic algorithm with 250 combinations tested) to maximize accuracy. The final LDA model was cross-validated with leave-one-out (LOO) cross-validation and the results used for performance metric calculation (confusion matrix, accuracy, ROC). Bootstrap ROC curves for the main model and comparative univariable LDA models were obtained by calculating the median of 100 bootstraps, as well as the boundaries where 75% of bootstrap curves lie.

