## Supplemental Information for "Iterative Extracellular Vesicle Protein Co-Expression Biomarker Refinement for Preoperative Classification of Histopathological Growth Patterns in Colorectal Liver Metastasis Patients"

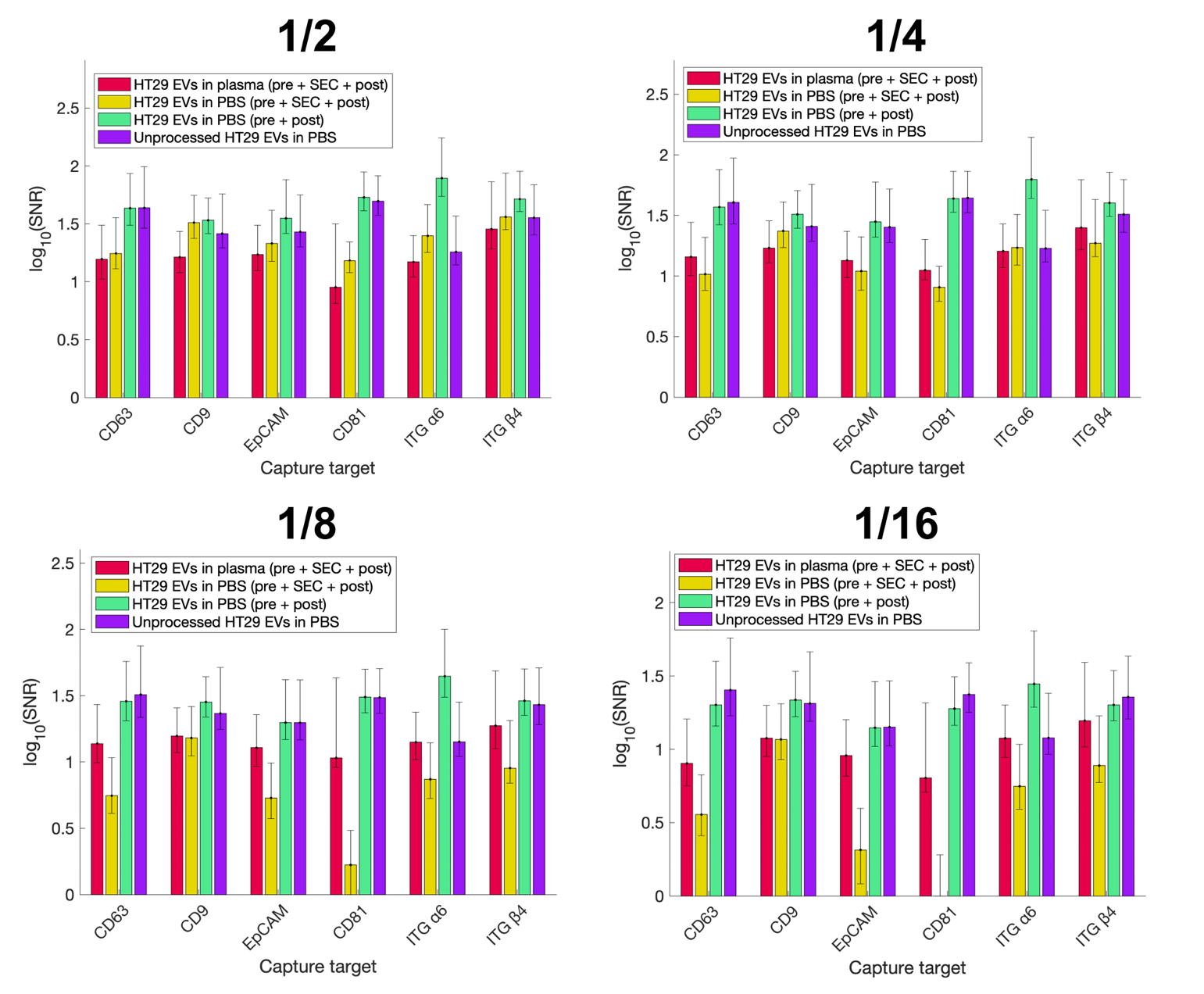

**Figure S1** Detection SNR of captured CFDA-SE-stained pre-purified HT29 EVs spiked 1/2-1/16 in plasma or PBS after each SEC processing step. The largest amount of signal loss occurs at the SEC purification step (compare green and yellow bars). Presence of a plasma matrix (red bars) is detrimental at low dilutions (1/2) but appears to have some blocking effect and prevent a portion of signal loss at higher dilutions (1/8, 1/16), even though signals remain lower than when the SEC step is omitted (green bars). Bars and error bars show the median and 95% confidence interval, respectively, obtained from bootstrapping technical replicates. Pre: centrifugation and filtration to remove debris prior to SEC; post: concentration of SEC-purified EVs.

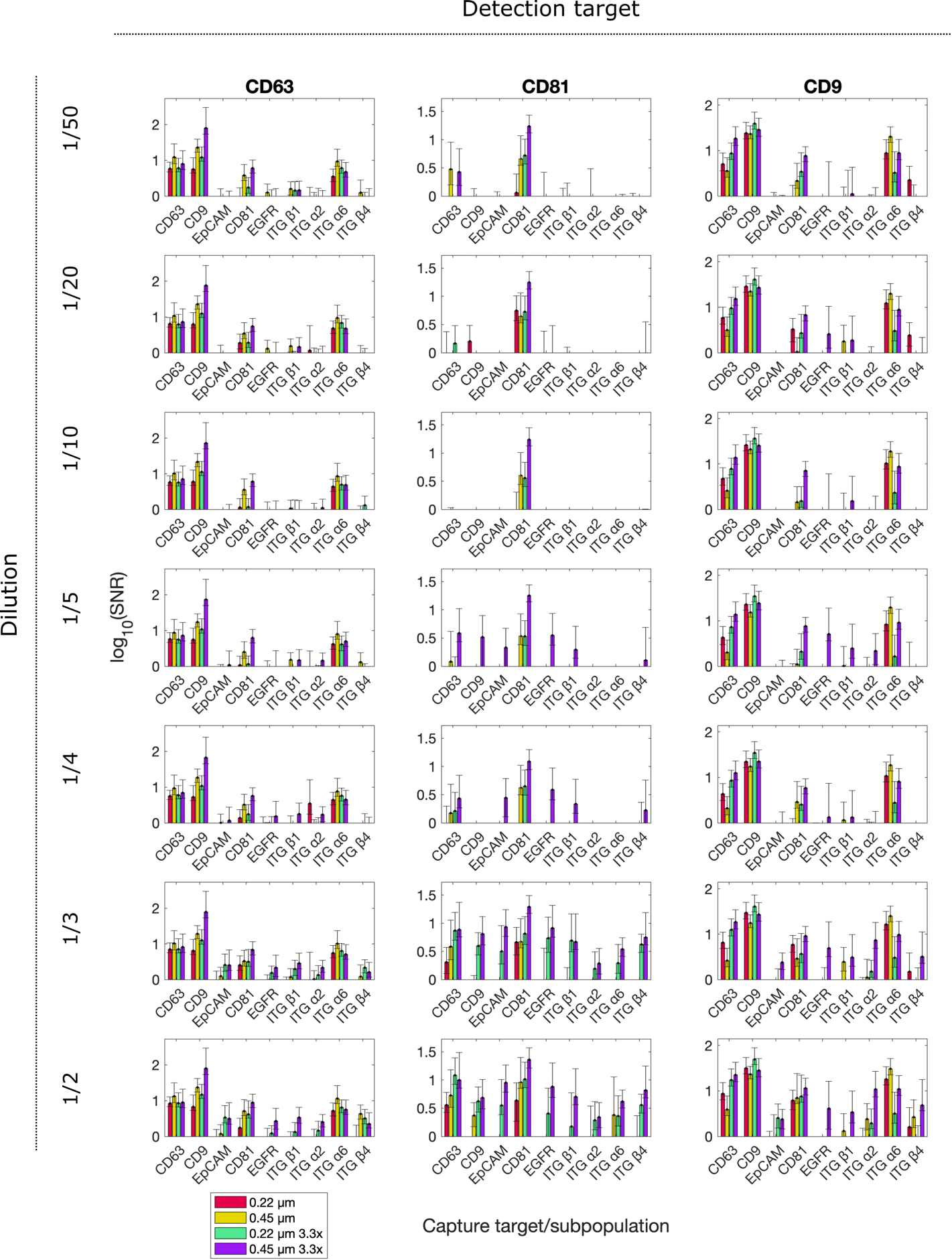

**Figure S2** Comparison of EVPio detection performance (log_10_[SNR]) of minimally purified pooled plasma-derived EVs when different filtration and concentration parameters are used. Filtration with 0.45 μm filters leads to better SNRs than 0.22 μm filtration. Pre-concentration of plasma modestly improves signal but increases non-specific binding and causes processing issues due to higher viscosity. Bars and error bars show the median and 95% confidence interval, respectively, obtained from bootstrapping technical replicates.

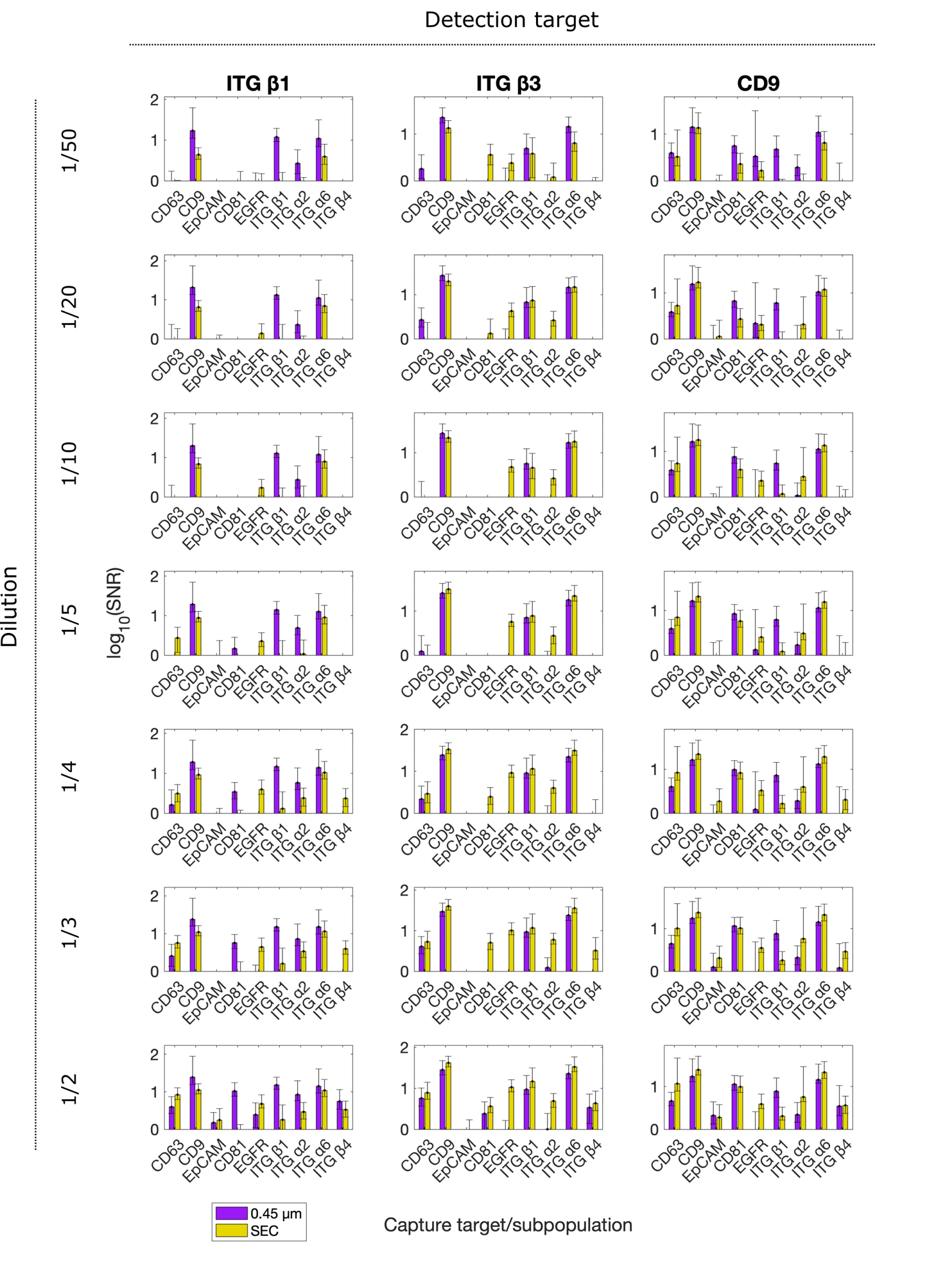

**Figure S3** Complete comparison of pooled plasma EVPio SNRs obtained at different dilution factors for minimally processed plasma (with 0.45 μm filtration) and SEC-processed plasma. Bars and error bars show the median and 95% confidence interval, respectively, obtained from bootstrapping technical replicates.

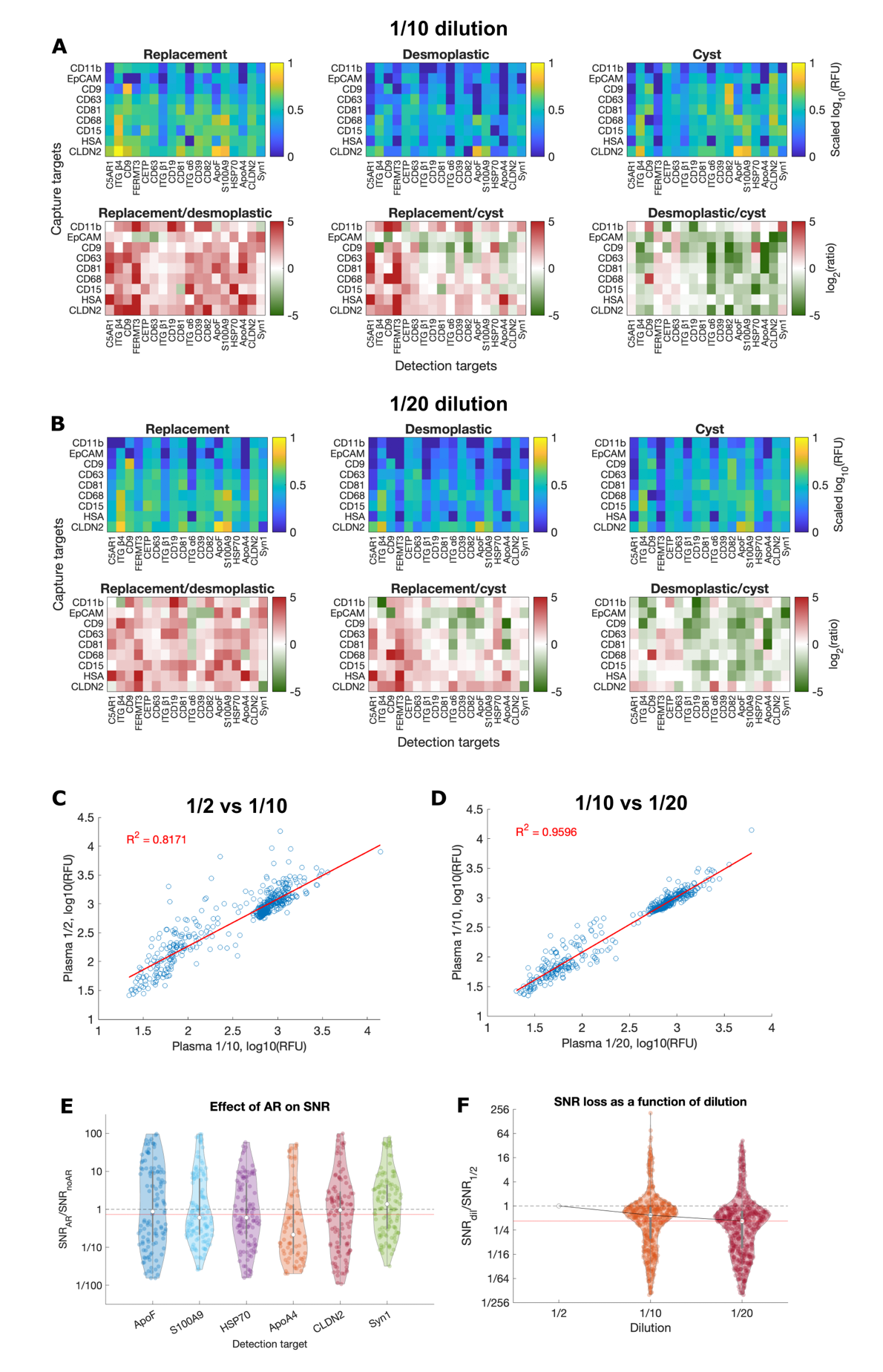

**Figure S4** Preliminary phenotyping of patient plasma EVs at (**A**) 1/10 and (**B**) 1/20 dilution in PBS and accompanying signal ratios maps between disease presentation (replacement, desmoplastic and cyst). (**C,D**) Raw RFU signals obtained at a given dilution strongly correlate with signals from other dilutions, supporting the reproducibility of EVPio-obtained phenotypes. (**E**) Ratio of signal obtained with and without AR for detection targets with known intravesicular localization. The absence of signal improveement with AR for most targets is likely due to the impact of freezing on EV integrity. The median SNR ratio is marked with a red line. (**F**) Median SNR drop with dilution factor, as a fraction of the SNR obtained at ½ dilution. The median SNR ratio at 1/20 dilution is marked as a red line. In violin plots, each dot represents the average of *n* = 10 technical replicates for a given (**E**) capture target and dilution or (**F**) co-expression pair.

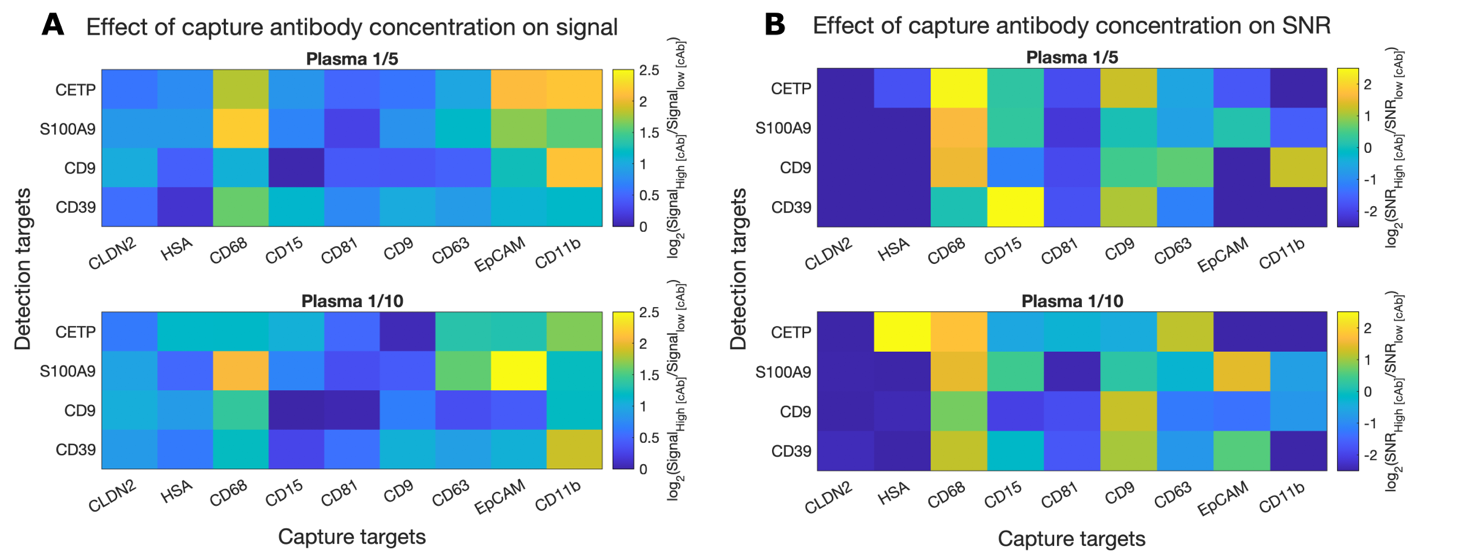

**Figure S5** Ratio of (**A**) signals and (**B**) SNRs obtained for capture antibodies patterned at 200-300 μg/mL and 100 μg/mL. A higher capture antibody concentration leads to increased signals for most co-expression pairs, with a median 1.9x signal boost (**A**). SNR improvements are more variable and depend on the expression level and level of antibody cross-reactivity of each pair (**B**).

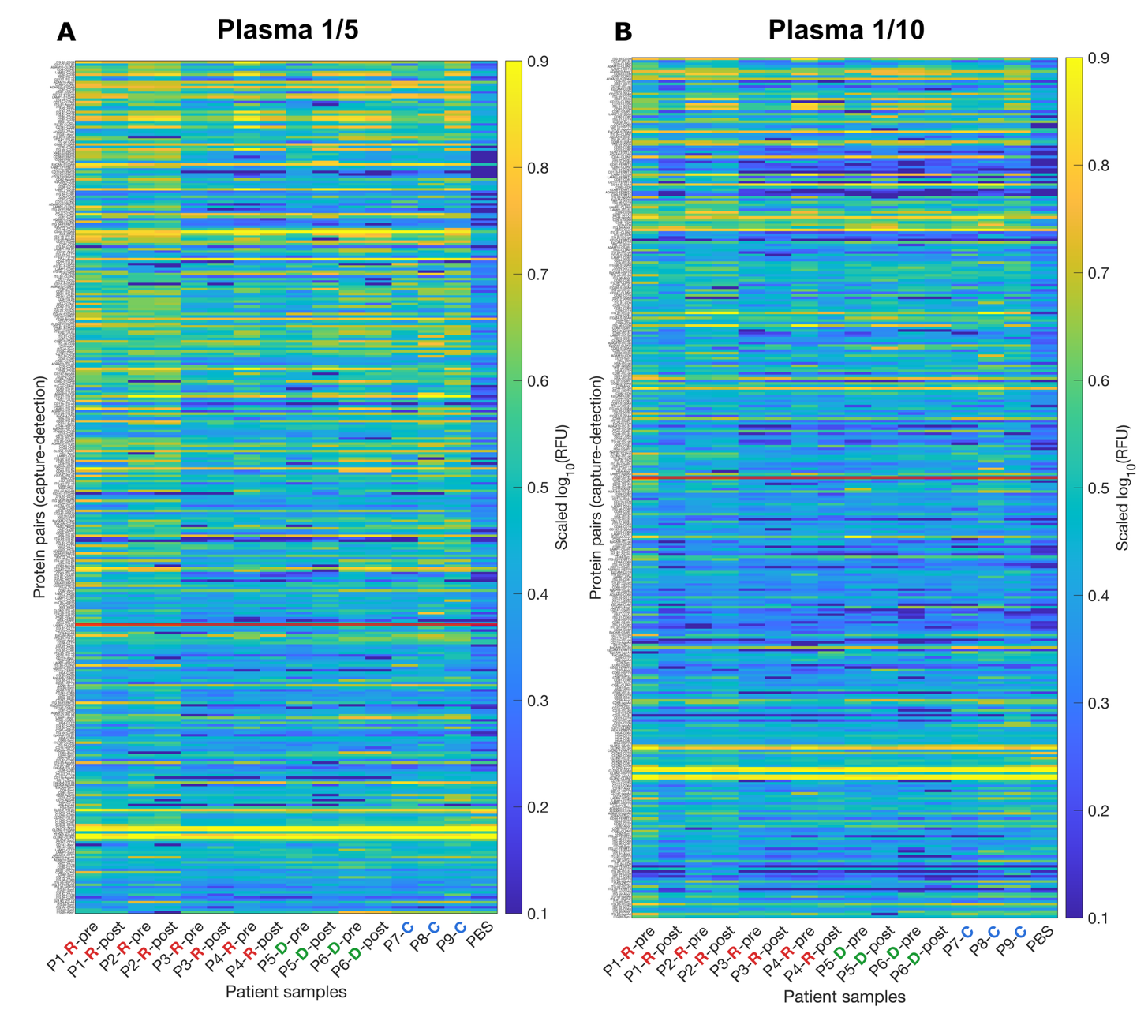

**Figure S6** Complete EVPio-CRCLM pilot phenotyping maps at (**A**) 1/5 and (**B**) 1/10 plasma dilution in PBS. Co-expression pairs on the vertical axis are ordered by decreasing SNR from the top, and the SNR cutoff of 3 is shown as a red horizontal line. **R**, replacement, **D**, desmoplastic, **C**, cyst, pre, before tumor resection, post, after tumor resection.

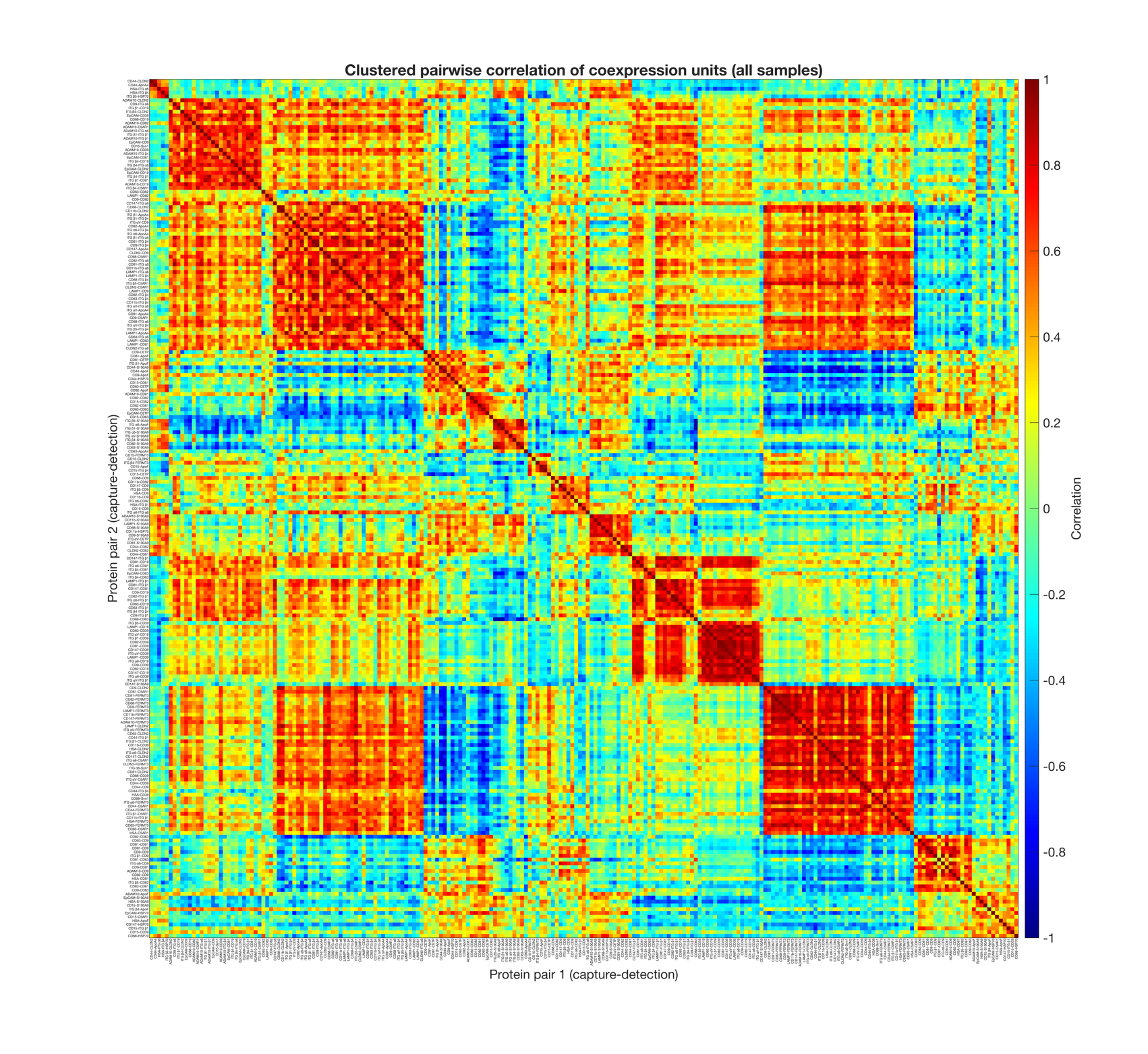

**Figure S7** Complete pairwise correlation map between all co-expression pairs measured in the EVPio-CRCLM pilot.

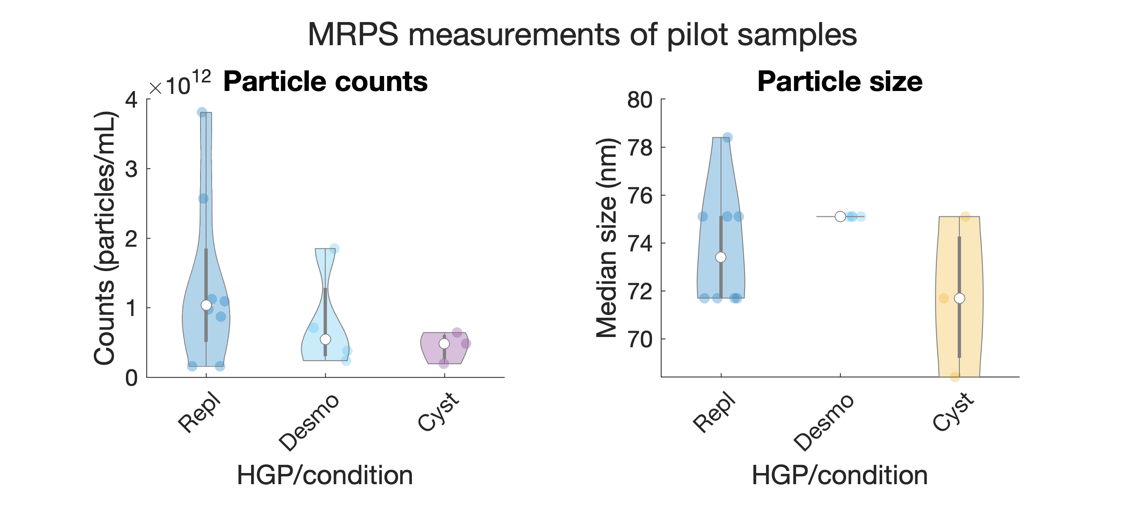

**Figure S8** Particle counts (*left*) and size (*right*) measure with microfluidic resistive pulse sensing (MRPS) of all samples included in the pilot (*n* = 9, with samples pre- and post-resection for CRCLM samples).

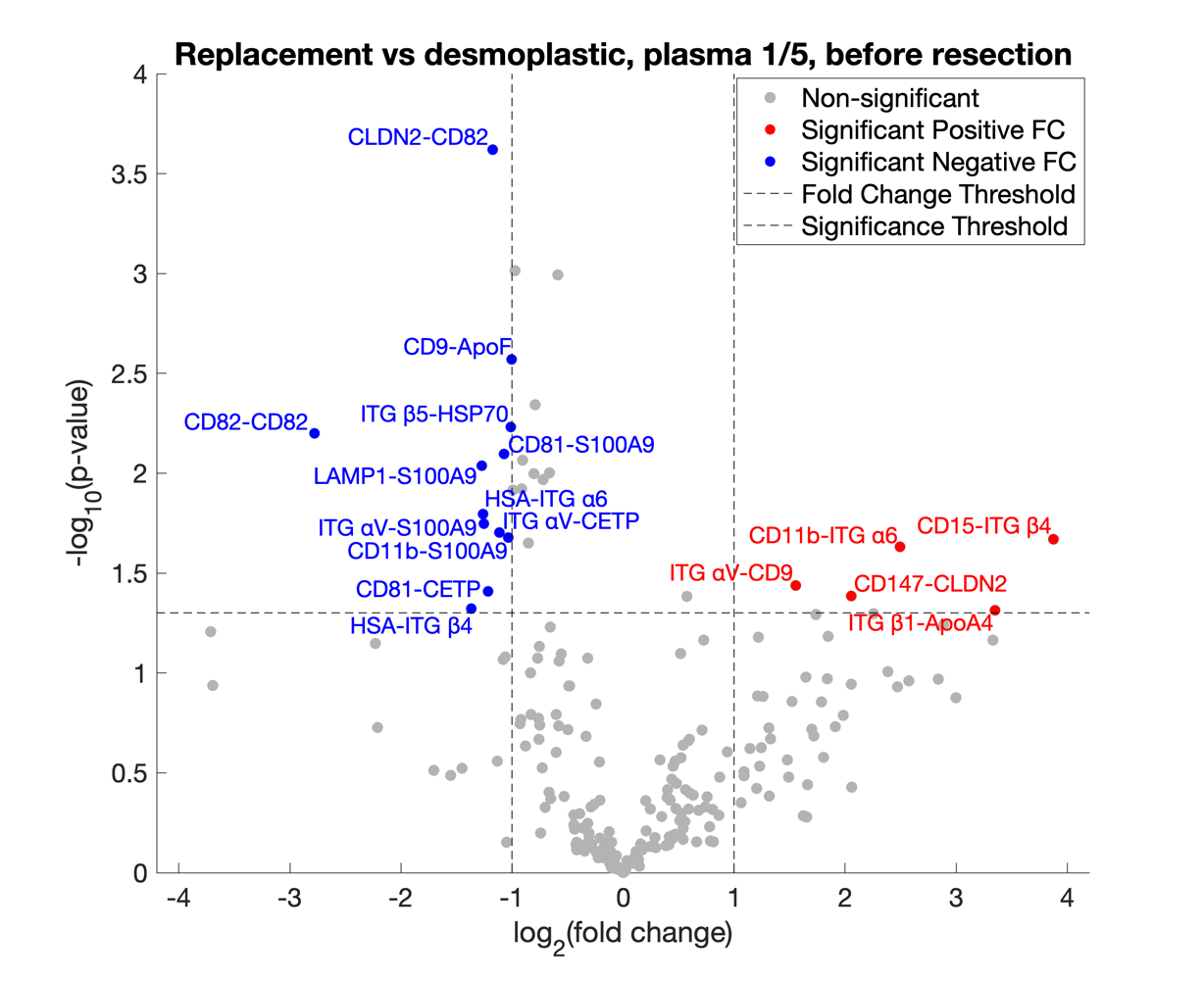

**Figure S9** Volcano plot showing the best discriminating co-expresion pairs from the EVPio-CRCLM pilot, for pre-resection, chemonaïve plasma samples diluted 1/5 in PBS. Combinations with notably higher signal in replacement and desmoplastic patients (fold change threshold of 2, significance threshold *p*-value = 0.05) are shown in red and blue, respectively.

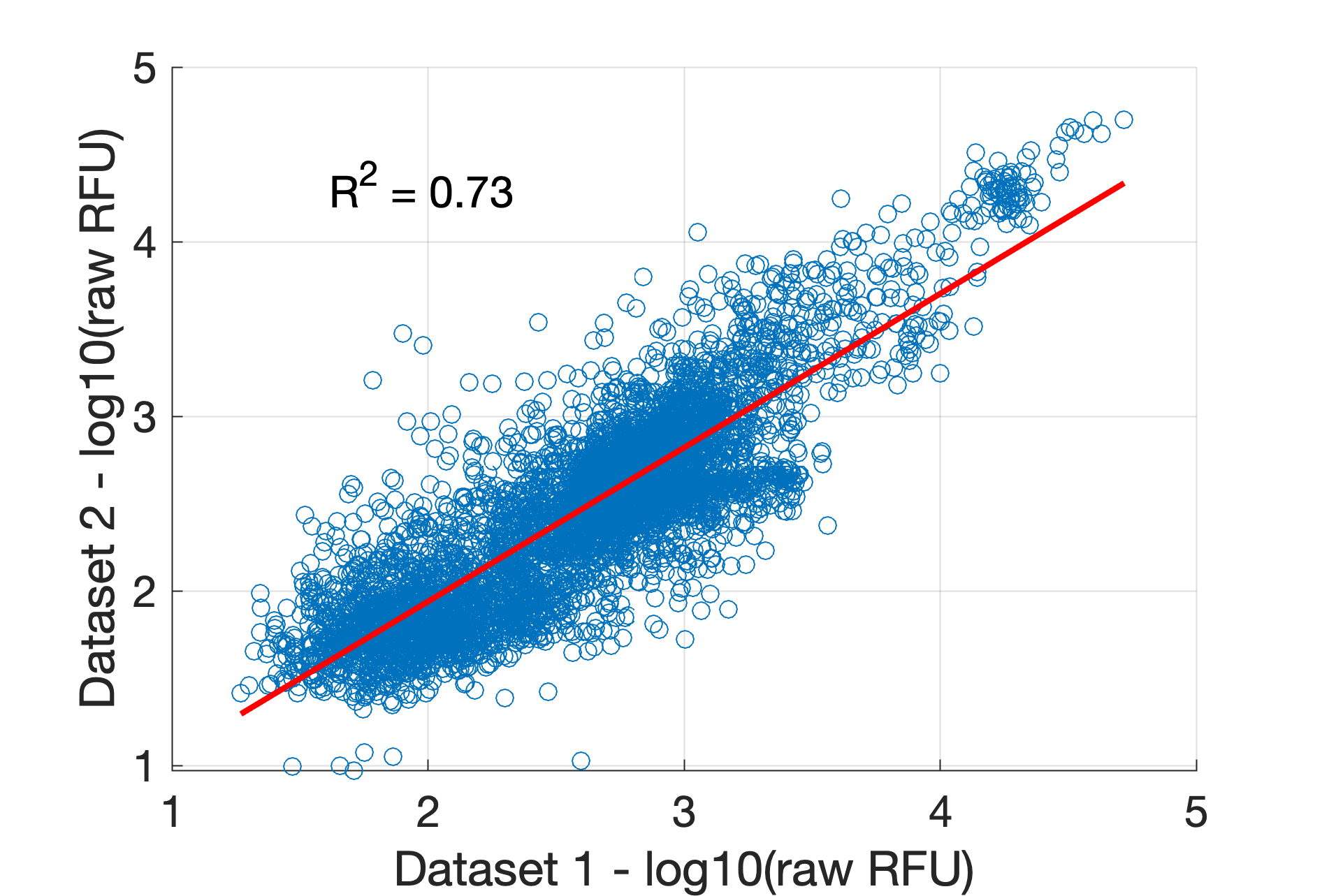

**Figure S10** Raw RFU signals obtained for each of the two independent EVPio-CRCLM assay runs show a strong positive correlation (R^2^ = 0.73), supporting assay reproducibility.

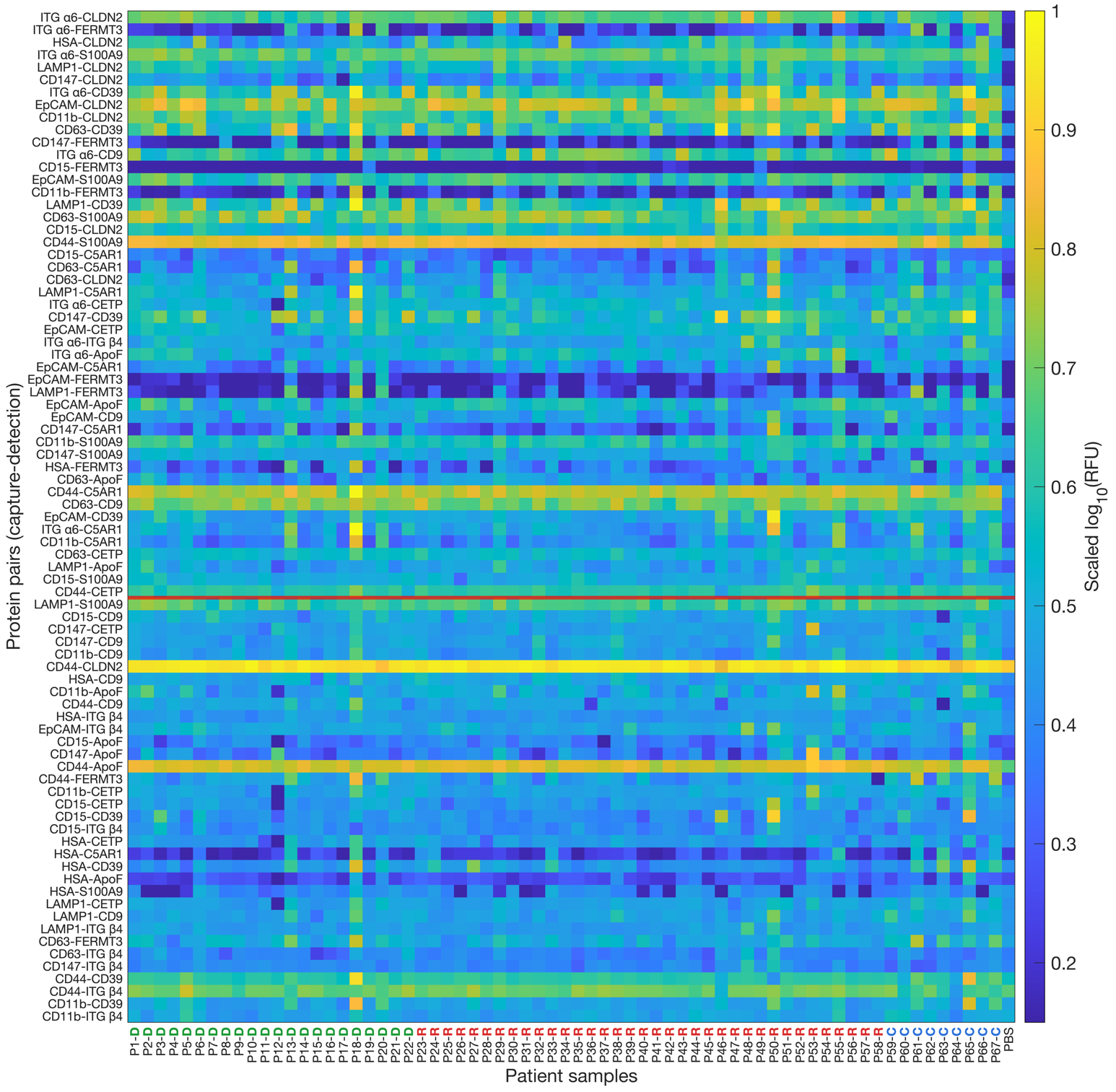

**Figure S11** Complete EVPio-CRCLM phenotyping heatmap of 58 CRCLM and 9 cyst patient plasma samples obtained from the batch-corrected combination of two independent assay runs (total of *n* = 20 experimental repeats except for combinations including CLDN2 and S100A9 for which *n* = 10). Protein co-expression pairs are sorted by decreasing SNR, and the SNR > 1.75 (92% confidence interval) criterium used for filtering out low-quality variables is shown as a red horizontal line. **D**, desmoplastic, **R**, replacement, **C**, cyst.

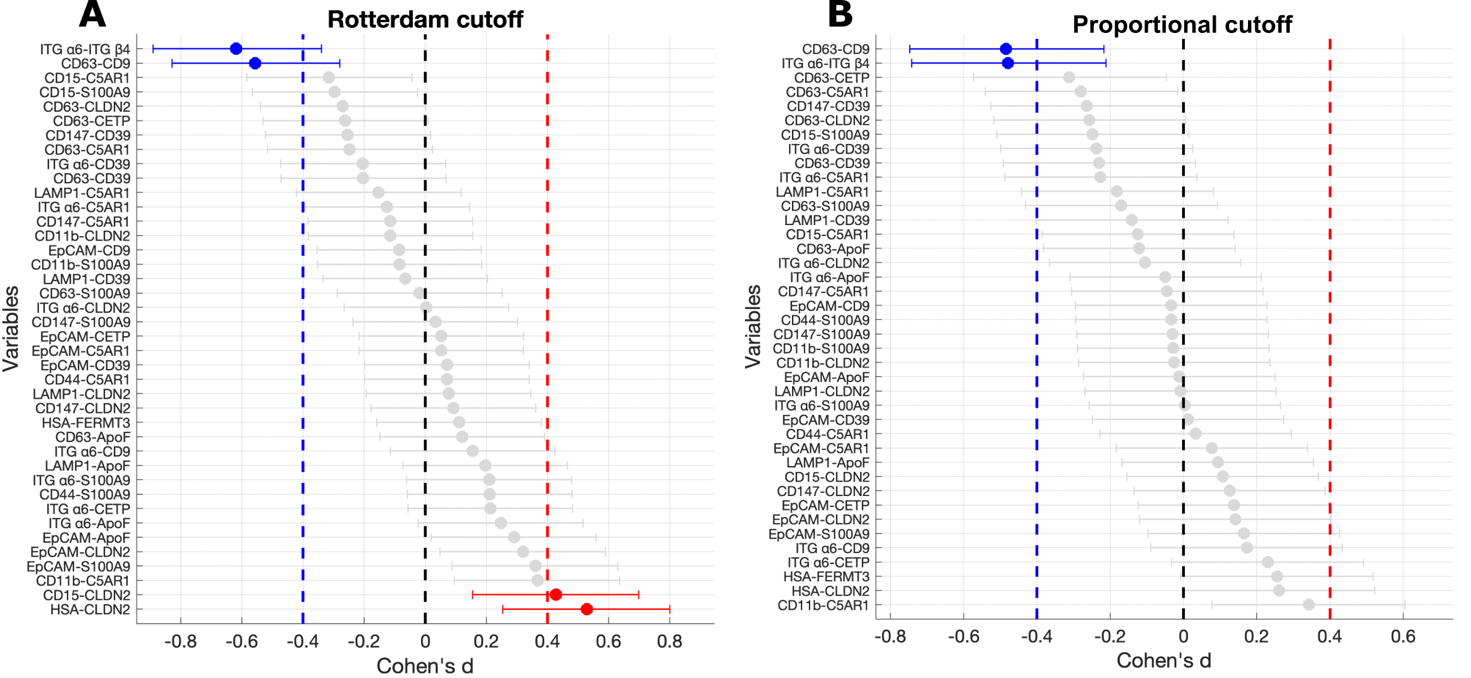

**Figure S12** Standardized effect size plots of quality-thresholded EVPio-CRCLM co-expression pairs for (**A**) Rotterdam cutoff- and (**B**) proportional (50%) cutoff-determined HGP classes. Combinations with an absolute Cohen’s *d* value above 0.4 are colored in red and blue for their stronger signal in replacement and desmoplastic samples, respectively.

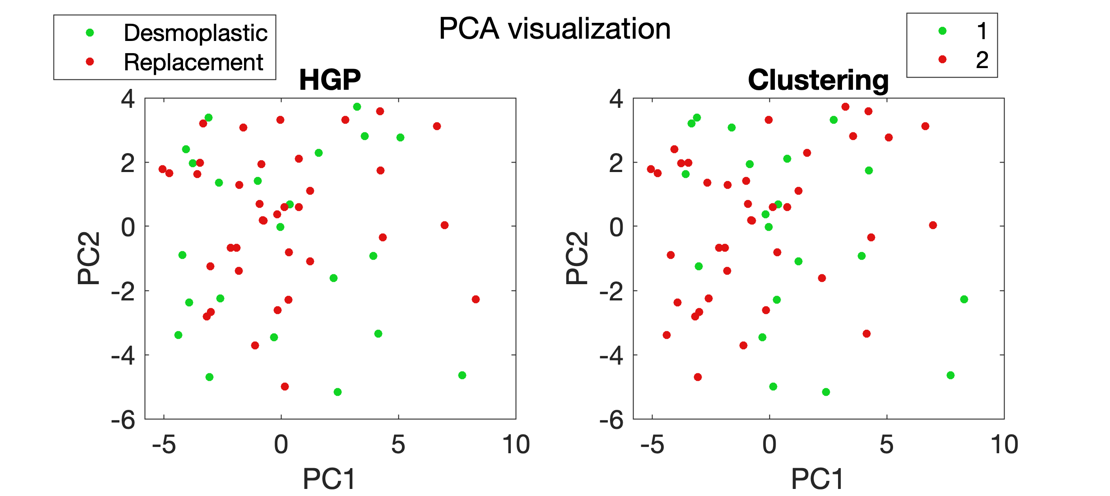

**Figure S13** Visualization of the principal component analysis (PCA) of the quality-filtered dataset with *p* = 40 features and *n* = 58 cancer patient samples and with superimposed (*left*) HGP labels and (*right*) gaussian mixture model-based clustering.

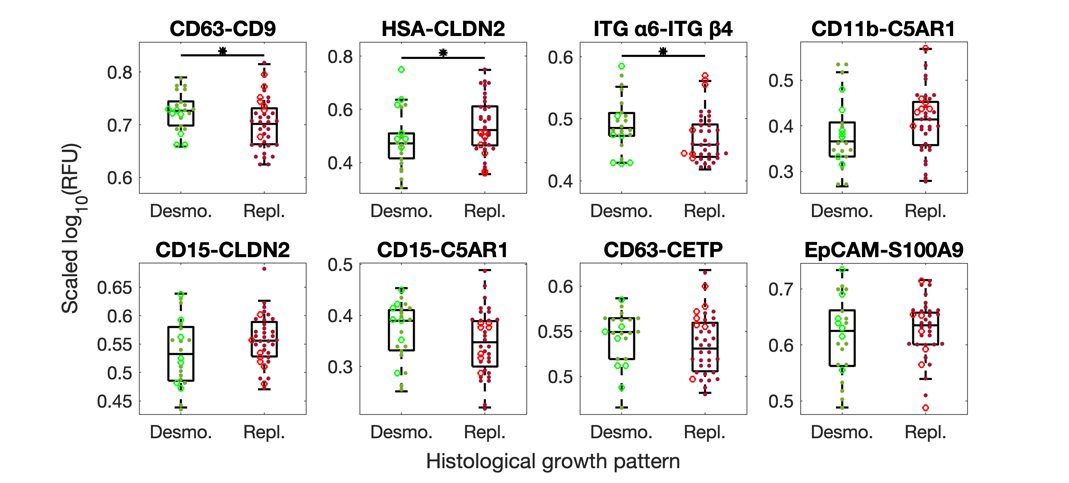

**Figure S14** Misclassification patterns within candidate co-expression pairs retained after ROC-based selection. Misclassified replacement samples (bright red open circles) tend to display higher CD63-CD9 signals (linked to stromal crosstalk), while misclassified desmoplastic samples (bright green open circles) exhibit higher than average HSA-CLDN2 signals (linked to hepatocyte mimicry), suggesting biological overlap or transitional states between HGPs. Significance threshold of *p*-value = 0.08.

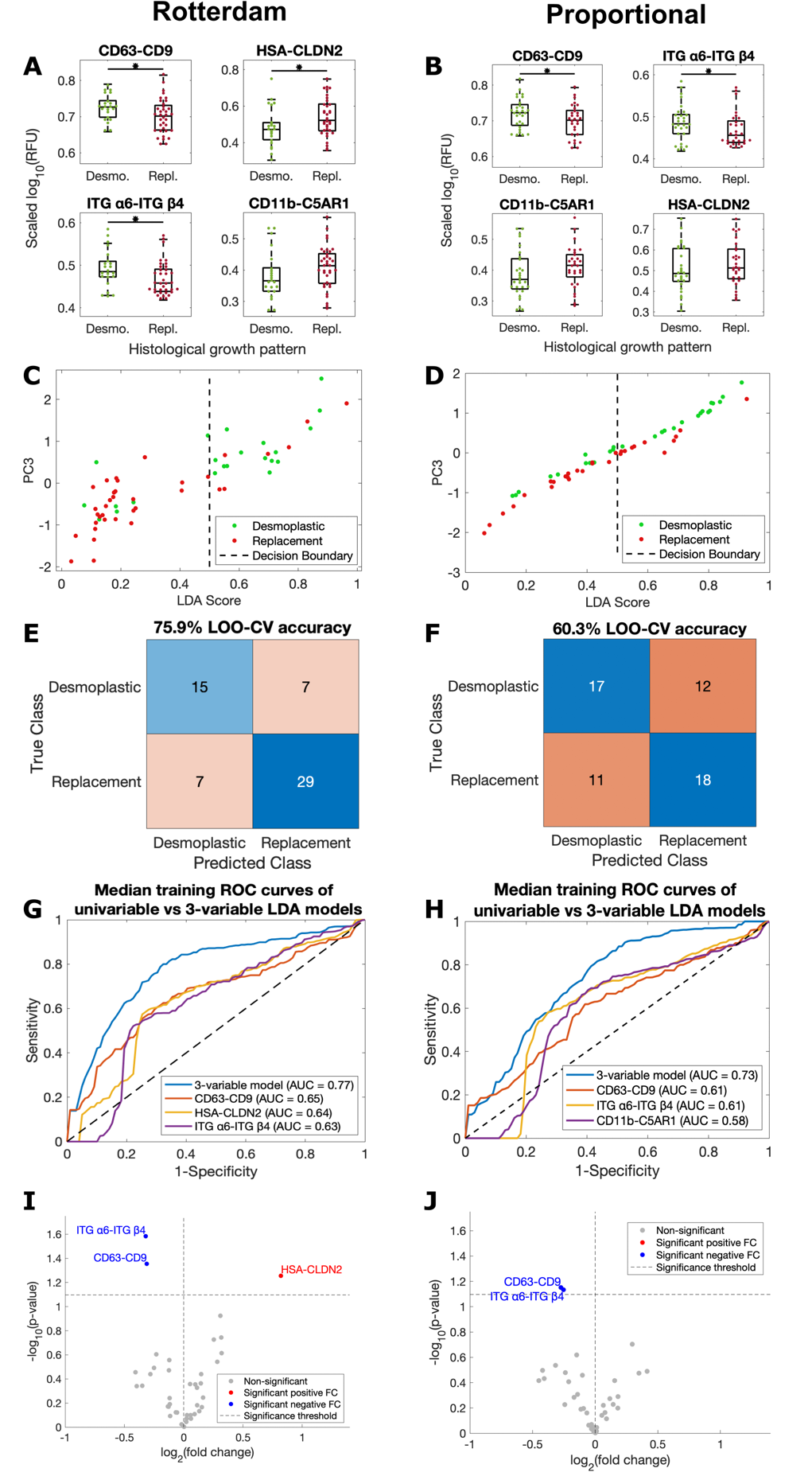

**Figure S15** Comparison of modeling performance when HGP class membership is defined according to the Rotterdam (*left*) or proportional (50%) (*right*) cutoff based on single-marker distributions (**A-B**), LDA axis separation (**C-D**), LDA classification accuracy (**E-F**), ROC analysis (**G-H**) and volcano plot analysis (threshold *p* < 0.08) of noise-filtered features (**I-J**). HSA-CLDN2 emerges as a driving, powerful composite predictor with Rotterdam-based patient classification.

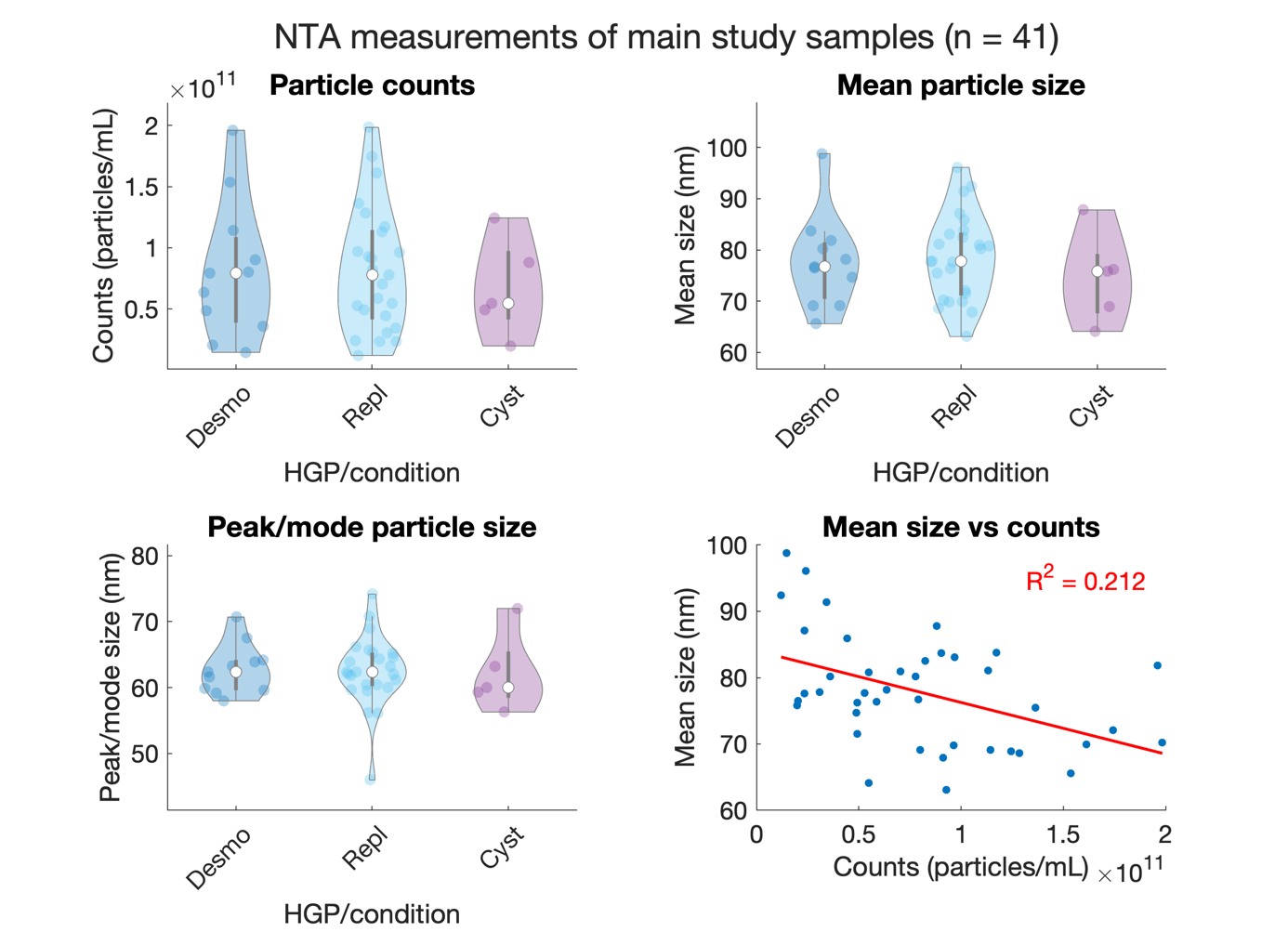

**Figure S16** Particle counts (*upper left*), mean size (*upper right*), and peak size (*lower left*) measured for a representative, randomly selected subset of samples from the main EVPio-CRCLM study (total *n* = 67). There is a weak negative correlation (R^2^ = 0.212) between mean size and counts (*lower right*).

**Table S1** Protein targets and antibodies used in the 4 main arms of this study

| **Antibody target** | **Capture/ Detection** | **Host species** | **Supplier** | **Catalog #** | **Lot #** | **EVPio assays** |
| --- | --- | --- | --- | --- | --- | --- |
| CD63 | Capture & detection | Mouse monoclonal | Biolegend | 353039 | B302182 | Optimization, preliminary, pilot |
| CD9 | Capture & detection | Mouse monoclonal | Biolegend | 312102 | B279344 | Optimization, preliminary, pilot |
| CD81 | Capture & detection | Mouse monoclonal | Biolegend | 349502 | B264968 | Optimization, preliminary, pilot |
| EpCAM | Capture | Mouse monoclonal | R&D Systems | MAB9601 | UTT1522011 | Optimization, preliminary, pilot, discovery |
| EGFR | Capture | Goat polyclonal | R&D Systems | AF231 | AUC1218041 | Optimization |
| Integrin β1 | Capture | Goat polyclonal | R&D Systems | AF1778 | WGJ0418111 | Optimization |
| Integrin α2 | Capture | Rat monoclonal | R&D Systems | MAB12332 | CBDM0118021 | Optimization |
| Integrin α6 | Capture | Mouse monoclonal | R&D Systems | MAB1350 | HNO052010A | Optimization |
| Integrin β4 | Capture | Mouse monoclonal | R&D Systems | MAB4060 | ZNC0219021 | Optimization, pilot |
| Integrin β1 | Detection | Mouse monoclonal | R&D Systems | MAB17781 | UUY0420081 | Optimization, preliminary |
| Integrin β3 | Detection | Mouse monoclonal | R&D Systems | MAB2266 | KUT0320081 | Optimization |
| CD11b | Capture | Rat monoclonal | Biolegend | 101202 | B349121 | Preliminary, pilot, discovery |
| CD68 | Capture | Mouse monoclonal | Biolegend | 333802 | B321990 | Preliminary, pilot |
| CD15 | Capture | Mouse monoclonal | Biolegend | 301902 | B349192 | Preliminary, pilot, discovery |
| Hepatocyte Specific Antigen (HSA) | Capture | Mouse monoclonal | Abcam | AB191200 | AB191200 | Preliminary, pilot, discovery |
| Claudin-2 | Capture | Mouse monoclonal | Sigma Aldrich | WH0009075M1 | H7251-3F1 | Preliminary, pilot |
| C5AR1 | Detection | Mouse monoclonal | Biolegend | 344302 | B295543 | Preliminary, pilot, discovery |
| Integrin β4 | Detection | Mouse monoclonal | R&D Systems | MAB4060 | ZNC0320011 | Preliminary |
| CD9 | Detection | Mouse monoclonal | Biolegend | 312102 | B353912 | Preliminary |
| FERMT3 | Detection | Sheep polyclonal | R&D Systems | AF7004 | CETM0212021 | Preliminary, pilot, discovery |
| CETP | Detection | Rabbit polyclonal | Invitrogen | PA5-89297 | YC3851329C | Preliminary, pilot, discovery |
| CD63 | Detection | Mouse monoclonal | Biolegend | 353014 | B240133 | Preliminary, pilot |
| CD19 | Detection | Mouse monoclonal | Biolegend | 302202 | B357499 | Preliminary, pilot |
| CD81 | Detection | Mouse monoclonal | Biolegend | 349502 | B342078 | Preliminary |
| Integrin α6 | Detection | Mouse monoclonal | R&D Systems | MAB1350 | HNO0520101 | Preliminary |
| CD39 | Detection | Mouse monoclonal | Biolegend | 328202 | B342001 | Preliminary, pilot, discovery |
| CD82 | Detection | Mouse monoclonal | R&D Systems | MAB4616 | ZVV0415121 | Preliminary |
| ApoF | Detection | Rabbit polyclonal | Invitrogen | PA5-118959 | YB3851081 | Preliminary, pilot |
| S100A9 | Detection | Mouse monoclonal | Biolegend | 600302 | B250234 | Preliminary, pilot, discovery |
| HSP70 | Detection | Rabbit polyclonal | Invitrogen | PA5-34772 | WD3258269B | Preliminary |
| ApoA4 | Detection | Mouse monoclonal | R&D Systems | MAB8125 | CLJV0219011 | Preliminary, pilot |
| Claudin-2 | Detection | Mouse monoclonal | Invitrogen | 32-5600 | WA317205 | Preliminary |
| Syntenin-1 | Detection | Mouse monoclonal | Invitrogen | MA5-32932 | YC3852151A | Preliminary |
| CD147 | Capture | Mouse monoclonal | Biolegend | 306202 | B262608 | Pilot, discovery |
| LAMP1 | Capture | Mouse monoclonal | Biolegend | 328602 | B366026 | Pilot |
| ADAM10 | Capture | Mouse monoclonal | R&D Systems | MAB1427 | HZR051710B | Pilot |
| CD44 | Capture | Rat monoclonal | Biolegend | 103014 | B269039 | Pilot, discovery |
| CD82 | Capture | Mouse monoclonal | R&D Systems | MAB4616 | ZVV061710B | Pilot |
| Integrin α6 | Capture | Mouse monoclonal | R&D Systems | MAB1350 | HNO0523011 | Pilot, discovery |
| Integrin β1 | Capture | Mouse monoclonal | R&D Systems | MAB17781 | UUY0316112 | Pilot |
| Integrin αv | Capture | Mouse monoclonal | R&D Systems | MAB1219 | HJH0218071 | Pilot |
| Integrin β5 | Capture | Mouse monoclonal | Invitrogen | 14-0497-82 | 2043810 | Pilot |
| CD9 | Detection | Mouse monoclonal | Biolegend | 312102 | B372349 | Pilot, discovery |
| CD81 | Detection | Mouse monoclonal | Biolegend | 349501 | B372461 | Pilot |
| Integrin β4 | Detection | Mouse monoclonal | R&D Systems | MAB4060 | ZNC0323031 | Pilot |
| Integrin β1 | Detection | Mouse monoclonal | R&D Systems | MAB17781 | UUY0321111 | Pilot |
| Integrin α6 | Detection | Mouse monoclonal | R&D Systems | MAB1350 | HNO0521101 | Pilot |
| CD82 | Detection | Mouse monoclonal | Biolegend | 342102 | 342102 | Pilot |
| Syntenin-1 | Detection | Mouse monoclonal | Invitrogen | MA5-32932 | YF3956732 | Pilot |
| HSP70 | Detection | Rabbit polyclonal | Invitrogen | PA5-34772 | YF3952117D | Pilot |
| Claudin-2 | Detection | Mouse monoclonal | Invitrogen | 32-5600 | XJ361834 | Pilot |
| LAMP1 | Capture | Mouse monoclonal | Biolegend | 328602 | B395496 | Discovery |
| CD63 | Capture | Mouse monoclonal | Biolegend | 353014 | B240133 | Discovery |
| Integrin β4 | Detection | Mouse monoclonal | R&D Systems | MAB4060 | ZNC0423111 | Discovery |
| ApoF | Detection | Rabbit polyclonal | Invitrogen | PA5-118959 | ZF4352482 | Discovery |
| Claudin-2 | Detection | Mouse monoclonal | Invitrogen | 32-5600 | ZD381741 | Discovery |

**Table S2** Sequences of the 15-mer barcodes used for EVPio assays.

| **Barcode** | **Sequence (5’→ 3’)** | **Antibody target** | **EVPio assay(s)** |
| --- | --- | --- | --- |
| BC03 | ATAAACTCGTCCAAT | FERMT3 | Preliminary |
|  |  |  | Pilot |
|  |  |  | Discovery |
| BC09 | ATGCTTCTACTCACT | ITG α6 | Preliminary |
|  |  |  | Pilot |
| BC23 | TAACACAAGCCAATG | HSP70 | Preliminary |
|  |  |  | Pilot |
| BC27 | TTACGACCTACAATG | ITG β1 | Optimization (comparison) |
|  |  |  | Preliminary |
|  |  |  | Pilot |
| BC32 | TAGCCTTTATTTCCA | CD9 | Optimization (comparison) |
|  |  |  | Preliminary |
|  |  |  | Pilot |
|  |  |  | Discovery |
| BC34 | TACCGTCCATTTCAA | ApoF | Preliminary |
|  |  |  | Pilot |
|  |  |  | Discovery |
| BC38 | TAGCCACTCATACGC | CD63 | Optimization |
|  |  |  | Preliminary |
|  |  |  | Pilot |
| BC43 | TCCATACAACATCAG | CD82 | Preliminary |
|  |  |  | Pilot |
| BC45 | TATCTCCCTTTCTTA | Syntenin-1 | Preliminary |
|  |  |  | Pilot |
| BC47 | AAAACGAAATCTCTG | CD39 | Preliminary |
|  |  |  | Pilot |
|  |  |  | Discovery |
| BC50 | TCAGCACTTCCTTAC | CD81 | Optimization |
|  |  |  | Preliminary |
|  |  |  | Pilot |
| BC51 | CGTCATTCACTACCT | CD9 | Optimization |
|  |  | S100A9 | Preliminary |
|  |  |  | Pilot |
|  |  |  | Discovery |
| BC53 | CGAACATCTAAACTA | CETP | Preliminary |
|  |  |  | Pilot |
|  |  |  | Discovery |
| BC61 | TAACATCCCTCATAG | ITG β3 | Optimization (comparison) |
|  |  | Claudin-2 | Preliminary |
|  |  |  | Pilot |
| BC67 | TTTCTTCACACGCTC | C5AR1 | Preliminary |
|  |  |  | Pilot |
|  |  |  | Discovery |
| BC71 | CTTTTCTACCCTATT | ApoA4 | Preliminary |
|  |  |  | Pilot |
|  |  |  | Discovery |
| BC75 | TACTAAGCCATCGTC | CD19 | Preliminary |
|  |  |  | Pilot |
| BC79 | CGATTCCTAACTATT | ITG β4 | Preliminary |
|  |  |  | Pilot |
|  |  |  | Discovery |

**Table S3** Aggregated clinical information for the patients included in the pilot and discovery arms of the EVPio-CRCLM study. Samples used in preliminary testing were drawn from the pilot cohort (for CRCLM) and the discovery cohort (for cyst).

| **EVPio assay** | **HGP/**  **condition** | | **Sex (M/F)** | **Age ± SD** | **# of lesions ± SD** | **HGP % ± SD (interface-weighted)** |
| --- | --- | --- | --- | --- | --- | --- |
| Pilot | Replacement (*n* = 4) | | 2/2 | 56 ± 11 | 1.8 ± 1.0 | 85 ± 19 |
|  | Desmoplastic (*n* = 2) | | 2/0 | 68 ± 18 | 1.0 ± 0.0 | 98 ± 4 |
|  | Cyst (*n* = 3) | | 1/2 | 52 ± 11 | N/A | N/A |
| Main (discovery) study | Replacement (*n* = 36) | Pure (*n* = 14) | 8/6 | 66 ± 8 | 3.0 ± 2.7 | 100 ± 0 |
|  |  | Mixed (*n* = 22) | 12/10 | 60 ± 10 | 2.6 ± 1.6 | 64 ± 38 |
|  | Desmoplastic (*n* = 22) | | 11/11 | 65 ± 12 | 2.5 ± 2.4 | 100 ± 0 |
|  | Cyst (*n* = 9) | | 0/9 | 65 ± 6 | N/A | N/A |

**Table S4** Credit attribution of composite design elements from Figure 1.

| **Element** | **Original Name** | **Creator** | **Source** | **Modifications** |
| --- | --- | --- | --- | --- |
| Hepatocyte | Hepatocyte Single Cell Biology | ScienceFigures.org | https://sciencefigures.org/figure/6401/hepatocyte_single_cell_biology/ | None |
| Fibroblast | Cancer Associated Fibroblasts Caf Cell | ScienceFigures.org | https://sciencefigures.org/figure/11189/cancer_associated_fibroblasts_caf_cell/ | None |
| Cancer cell | Cancer Cell 2 | ScienceFigures.org | https://sciencefigures.org/figure/8865/cancer_cell_2/ | None |
| Blood vessel | Main Blood Vessel Ramifications | ScienceFigures.org | https://sciencefigures.org/figure/11802/main_blood_vessel_ramifications/ | Color |
| Human with arm blood vessels | Prostate Lung Liver Tumor | ScienceFigures.org | https://sciencefigures.org/figure/9823/prostate_lung_liver_tumor/ | Removed organs, kept outline and blood vessels |
| Liver & gastrointestinal tract | Digestive System Body | ScienceFigures.org | https://sciencefigures.org/figure/167/digestive_system_body/ | Removed outline and trachea/esophagus |
| Syringe | Syringe 2 Lateral | ScienceFigures.org | https://sciencefigures.org/figure/5854/seringue_2_lateral/ | None |
| Blood tube | Tube 2 | ScienceFigures.org | https://sciencefigures.org/figure/564/tube_2/ | Inner liquid color |
| Plasma separation tube | Leucosep Tube Pbmc Ring | ScienceFigures.org | https://sciencefigures.org/figure/12116/leucosep_tube_pbmc_ring/ | None |
| Cryo tube | Cryovial Flask Clamp Zoom | ScienceFigures.org | https://sciencefigures.org/figure/7274/cryovial_flask_clamp_zoom/ | Inner liquid color |
| Cold warning sign | Low Temperature Sign | ScienceFigures.org | https://sciencefigures.org/figure/3266/low_temperature_sign/ | None |
| Open centrifuge | Hand Eppendorf Centrifuge Lab Zoom | ScienceFigures.org | https://sciencefigures.org/figure/5567/hand_eppendorf_centrifuge_lab_zoom/ | None |
| Test tube | Microtube 2 | ScienceFigures.org | https://sciencefigures.org/figure/395/microtube_2/ | Inner liquid color |
| Extracellular vesicle | Hybrid Exosome | ScienceFigures.org | https://sciencefigures.org/figure/9811/hybrid_exosome/ | None |
| Sample rack | Vial Flask Hplc Autosampler Rack | ScienceFigures.org | https://sciencefigures.org/figure/5573/vial_flask_hplc_autosampler_rack/ | Tube colors & positions |
